# Functional alignment of protein language models via reinforcement learning

**DOI:** 10.1101/2025.05.02.651993

**Authors:** Nathaniel Blalock, Srinath Seshadri, Kensuke Nakamura, Agrim Babbar, Sarah A Fahlberg, Ameya Kulkarni, Philip A Romero

## Abstract

Protein language models (pLMs) enable generative design of novel protein sequences but remain fundamentally misaligned with protein engineering goals, as they lack explicit understanding of function and often fail to improve properties beyond those found in nature. We introduce Reinforcement Learning from eXperimental Feedback (RLXF), a general framework that aligns protein language models with experimentally measured functional objectives, drawing inspiration from the methods used to align large language models like ChatGPT. Applied across five diverse protein families, RLXF improves generation of high-functioning variants beyond pre-trained baselines. We demonstrate this with CreiLOV, an oxygen-independent fluorescent protein, where RLXF-aligned models generate sequences with significantly enhanced fluorescence, including the most fluorescent CreiLOV variants reported to date. Our results indicate that RLXF-aligned models effectively integrate the evolutionary knowledge encoded in pre-trained pLMs with experimental observations, improving the success rate of generated sequences and enabling the discovery of synergistic mutation combinations that are difficult to identify through zero-shot or evolutionary approaches. RLXF provides a scalable and accessible approach to steer generative models toward desired biochemical properties, enabling function-driven protein design beyond the limits of natural evolution.

## Introduction

Protein language models (pLMs) leverage advances in natural language processing to learn how a protein’s linear arrangement of amino acids encodes its structure, function, and dynamics. Built on transformer-based architectures, pLMs generate representations that capture the biological meaning of a protein such as its fold, molecular activity, or evolutionary role rather than relying solely on sequence similarity^1,2^3/6/26 3:51:00 PM. These learned representations have proven powerful across a range of tasks including structure prediction^3,4^3/6/26 3:51:00 PM, function annotation^5,6^3/6/26 3:51:00 PM, variant effect prediction^7,8^3/6/26 3:51:00 PM, and evolutionary modeling^9,10^3/6/26 3:51:00 PM. More recently, pLMs have emerged as state-of-the-art tools for generative protein design, enabling the creation of novel and functional sequences including variants of green fluorescent protein^1,11^3/6/26 3:51:00 PM, lysozyme^12^3/6/26 3:51:00 PM, carbonic anhydrases^13^3/6/26 3:51:00 PM, lactate dehydrogenases^13^3/6/26 3:51:00 PM, triosephosphate isomerases^14^3/6/26 3:51:00 PM, and peptide binders^15,16^3/6/26 3:51:00 PM that differ substantially from any known natural proteins.

pLMs such as UniRep^9^3/6/26 3:51:00 PM and Evolutionary Scale Modeling (ESM)^3^3/6/26 3:51:00 PM are trained on large collections of natural protein sequences sampled from the evolutionary tree. Using self-supervised learning objectives such as autoregressive next-token prediction^12,17–19^3/6/26 3:51:00 PM or masked language modeling^1,20–22^3/6/26 3:51:00 PM, these models learn context-aware representations that capture structural features, biological functions, and evolutionary constraints embedded in natural proteins. When deployed for sequence generation, pLMs produce highly diverse outputs that mirror the statistical patterns and correlations present in natural proteins^12^3/6/26 3:51:00 PM. Although these sequences may be far from any individual example in the training data, they are still drawn from the same distribution, leading to novelty in sequence but not in function. Because pLMs have no explicit understanding of function, they often fail to generate proteins with enhanced or non-natural activities^1,11,12^. As a result, they remain fundamentally misaligned with the core goal of protein engineering: to create proteins beyond what evolution has already explored.

In this work, we introduce Reinforcement Learning from eXperimental Feedback (RLXF), a framework for aligning pLMs with experimentally derived notions of biomolecular function. This enables generative design of proteins with enhanced properties tailored to user-defined objectives. Our approach draws inspiration from the reinforcement learning techniques that aligned large language models with human preferences, resulting in transformative tools such as ChatGPT^23^3/6/26 3:51:00 PM and Claude^24^3/6/26 3:51:00 PM. In RLXF, a reward function, such as a supervised sequence-function predictor or any sequence scoring model, provides feedback to the pLM, guiding it to generate sequences with improved function. The outcome is a functionally aligned model that can be repeatedly sampled to produce diverse sequences optimized for the desired property.

We apply RLXF across five diverse protein classes to demonstrate its generalizability and effectiveness at generating optimized sequences by learning functional constraints beyond those captured during pre-training. As a case study, we align the 650M parameter ESM-2 model to experimental fluorescence data from the CreiLOV flavin-binding fluorescent protein. The aligned model learns to prioritize mutations that enhance fluorescence, many of which are missed by the base model. Experimental validation reveals the RLXF-aligned model generates a higher fraction of functional sequences, a greater number of sequences more fluorescent than CreiLOV, and several variants more fluorescent than the brightest known flavin-binding fluorescent protein. Our results highlight the importance of aligning pLMs with engineering objectives to move beyond natural distributions and unlock new functional potential in protein design.

## Results

### Functional alignment of protein language models

Large language models (LLMs) are pre-trained with the sole task of text completion, learning the syntax and semantics of language, but not capturing human intent or preferences. The key breakthrough behind ChatGPT was the use of reinforcement learning from human feedback (RLHF) to align pre-trained LLMs with human goals, resulting in models that generate more useful, truthful, and safe responses^23–26^3/6/26 3:51:00 PM. Protein language models share many parallels with LLMs. They generate syntactically and semantically valid protein sequences, but these sequences are not necessarily useful for human biotechnology applications such as sustainable energy, chemistry, or medicine. pLMs must likewise be aligned with human intent to generate proteins with biophysical or biochemical properties specified by humans.

Inspired by RLHF, we introduce Reinforcement Learning from eXperimental Feedback (RLXF) for aligning pLMs to generate sequences with enhanced or non-natural activities (Fig. 1a). At the core of RLXF is a user-defined *reward model* that modifies the base pLM to generate sequences with high reward values. This reward model can be any sequence- or structure-based scoring function. We focus on supervised models trained on experimental sequence-function data because they accurately capture complex biophysical and biochemical properties^27–29^3/6/26 3:51:00 PM.

**Figure 1.**
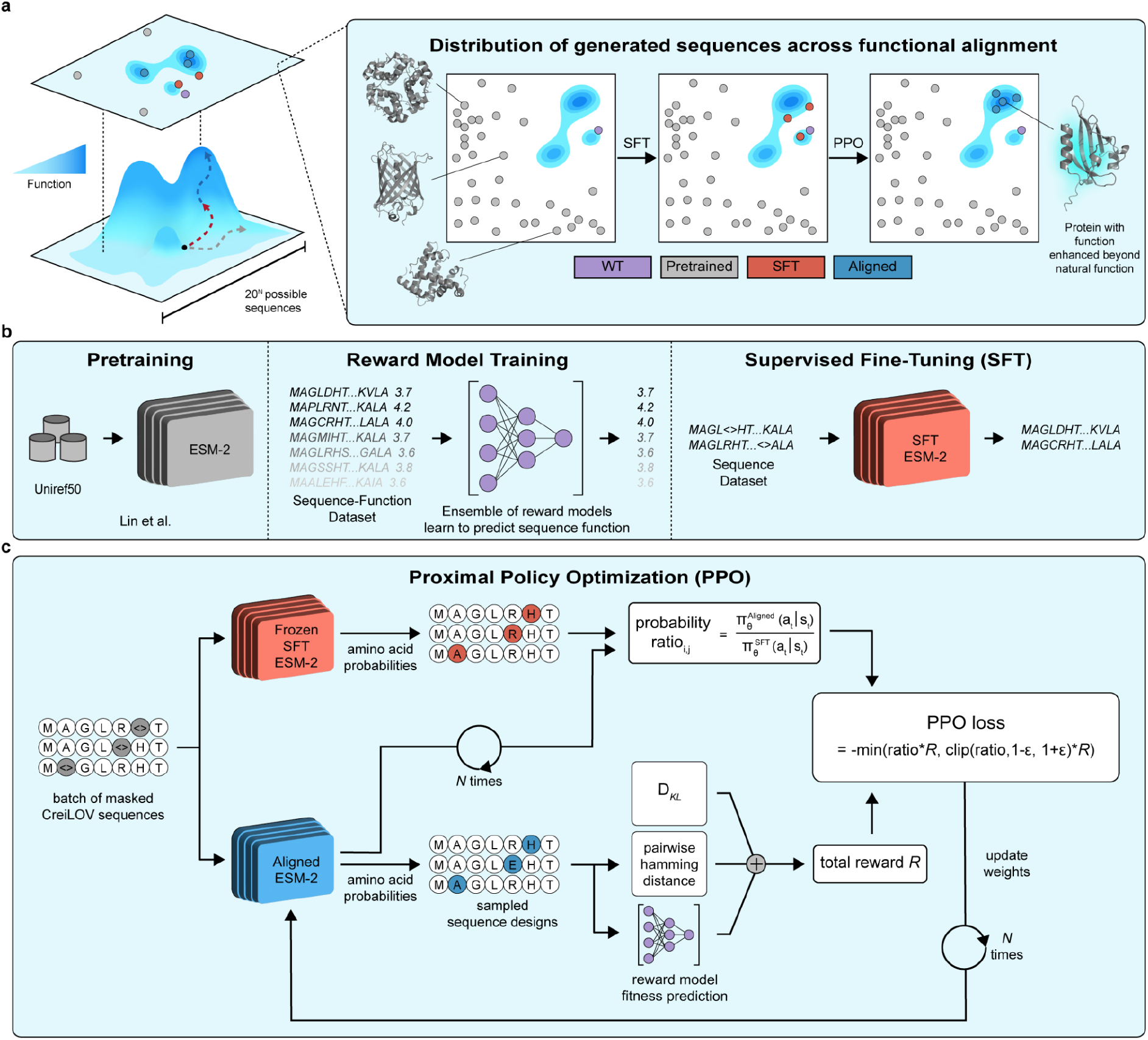
Reinforcement Learning from eXperimental Feedback (RLXF). **a**, RLXF aligns the generative distribution of protein language models with experimental measures of function, biasing sequence designs toward enhanced activity. **b**, RLXF begins with a pre-trained pLM such as ESM-2. A reward model is trained on sequence-function data and used to align the generative distribution of the pLM. The supervised fine-tuning (SFT) phase initializes the model in a functionally enriched region of sequence space, guiding it toward high-reward sequences. **c**, PPO directly optimizes the generative performance of the pLM. PPO iteratively updates ESM-2 weights *N* times each epoch to become more likely to generate high reward and diverse sequence designs. For each sampled amino acid, the PPO objective uses a ratio of probabilities: the numerator contains the probability assigned by the current (aligned) model, and the denominator contains the probability assigned by the previous (SFT) model to the same amino acid at that position. Gradients are backpropagated through the aligned model’s probabilities, allowing it to adjust its generative distribution toward higher-reward sequences. The total reward consists of three components: a normalized sequence-function reward model score, a pairwise Hamming distance term to encourage sequence diversity, and a Kullback-Leibler divergence penalty (*D*_*KL*_) to constrain updates and prevent forgetting of pre-trained knowledge in ESM-2. To stabilize training, the PPO loss is clipped using a threshold ε to limit the size of policy updates.

RLXF follows a two-phase training strategy analogous to RLHF: *supervised fine-tuning* (SFT) that helps initialize the model in the correct region of sequence space, followed by *proximal policy optimization* (PPO) that directly aligns sequence generation with the reward model (Fig. 1b-c). In the SFT phase, we use the reward model to explore sequence space and identify a collection of high-reward sequences. These are then used to fine-tune the generative pLM, biasing it to sample amino acids observed in these desirable sequences. In the PPO phase, sequences are sampled directly from the pLM, scored by the reward model, and the resulting reward values are fed back to update the pLM to further increase the likelihood of generating high-reward sequences. Together, SFT and PPO align the generative pLM with human-defined objectives, enabling it to produce protein sequences with optimized functional properties.

RLXF is a general framework that can be applied to any protein sequence generation model, including variational autoencoders^30^3/6/26 3:51:00 PM, structure-based graph neural networks like ProteinMPNN^31^3/6/26 3:51:00 PM, sequence diffusion models like EvoDiff^32^3/6/26 3:51:00 PM, autoregressive models like ProGen^2,12,33^3/6/26 3:51:00 PM, and masked language models like ESM-2^3^3/6/26 3:51:00 PM. In this work, we focus primarily on the ESM family of models using masked language modeling for sequence generation. We developed and evaluated RLXF using the CreiLOV flavin-binding fluorescent protein. We trained a supervised reward model with existing deep mutational scanning (DMS) data of CreiLOV variants^34^3/6/26 3:51:00 PM to predict fluorescence from sequence (Supplementary Figure 1). We used CreiLOV to systematically explore the generative capabilities of pre-trained ESM models, different generative sampling strategies, supervised fine-tuning approaches, parameter-efficient training techniques, ESM model scaling effects, and benchmark RLXF against the related method direct preference optimization (Supplementary Figure 1-7).

### Functional alignment improves sequence generation across diverse protein classes

Protein language models such as ESM-2 lack an explicit notion of biomolecular function and therefore struggle to generate proteins with enhanced or non-natural activities. RLXF provides a general framework for aligning pLMs with human-defined functional objectives, enabling the generation of sequences with improved and tunable biochemical properties. To evaluate the generality of RLXF, we applied it to five diverse protein classes spanning a range of sizes, structural folds, and molecular functions: poly(A)-binding protein (Pab1)^35^3/6/26 3:51:00 PM, β-glucosidase (Bgl3)^36^3/6/26 3:51:00 PM, ubiquitination factor E4B (Ube4b)^37^3/6/26 3:51:00 PM, *Aequorea victoria* green fluorescent protein (avGFP)^38^3/6/26 3:51:00 PM, and the protein G B1 domain (GB1)^39^3/6/26 3:51:00 PM (Supplementary Table 1; Supplementary Figure 8). We kept most hyperparameters consistent with those chosen during RLXF development on CreiLOV (Supplementary Table 2-9). For each system, we trained a supervised reward model on DMS data and used it to align ESM-2 models of increasing size (8M, 35M, 150M, and 650M parameters) (Supplementary Table 2-4).

To better understand how RLXF incorporates functional information to improve sequence generation, we compared three model variants: the base ESM-2 models, models trained only with supervised fine-tuning (SFT), and models trained with both SFT and PPO. We evaluated generation performance using a win rate metric commonly used in natural language processing (NLP)^23,26^3/6/26 3:51:00 PM, defined here as the percentage of times a model generates a sequence with a higher predicted reward than one generated by the base ESM-2 650M model. Across protein classes and model sizes, RLXF significantly increased the win rate compared to the base model (Fig. 2a). No single model size consistently outperformed the others. The best-performing model varied depending on the specific protein system.

**Figure 2.**
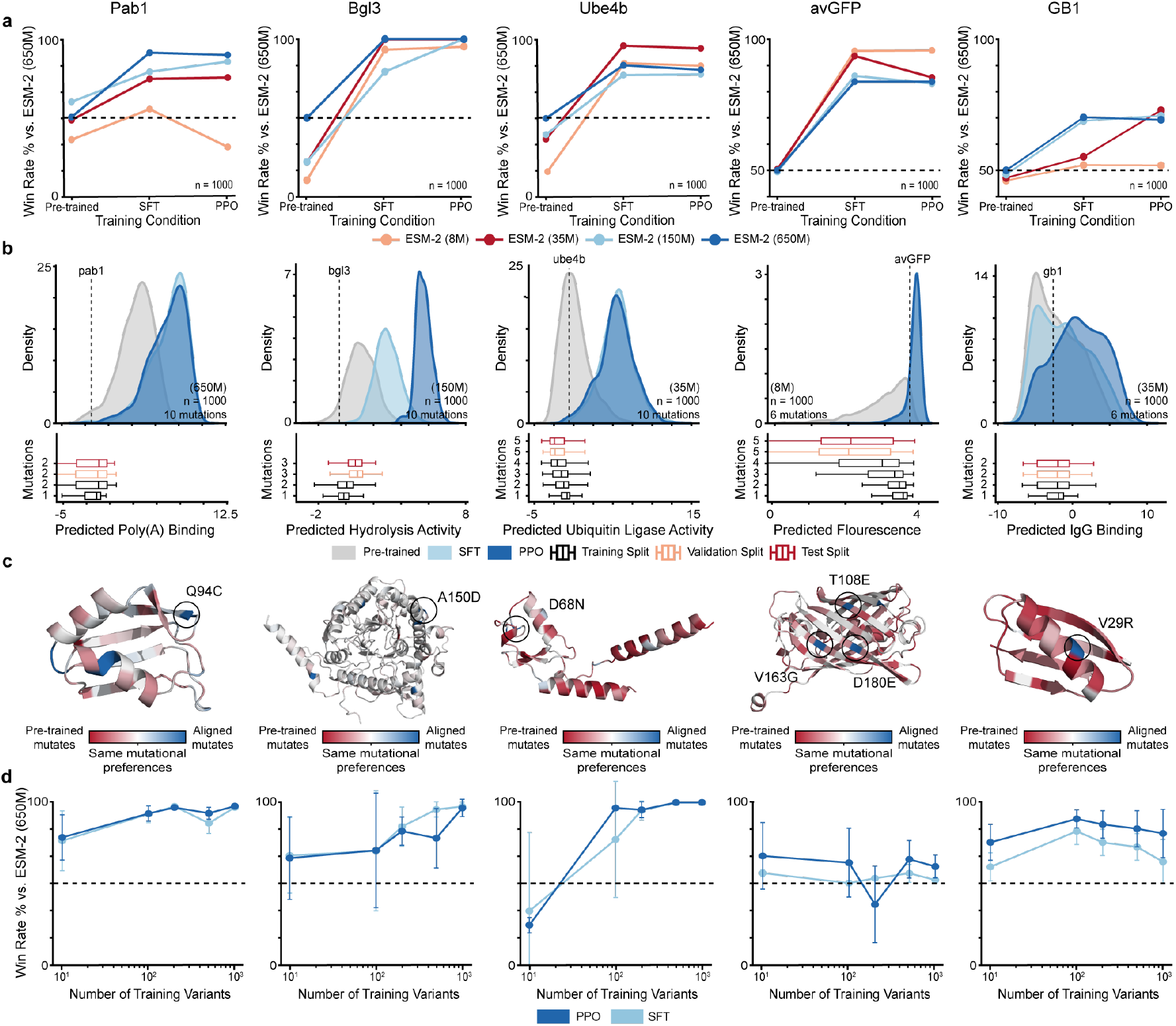
Functional alignment improves properties of generated sequences. **a**, Win rate of sequence designs from pre-trained, SFT, and PPO ESM-2 models of varying sizes relative to pre-trained ESM-2 (650M). **b**, Kernel density estimate plots depicting the distribution of predicted function for 1,000 designs generated by ESM-2 at each stage of functional alignment and boxplots for predicted function of sequences in reward model training, validation, and test splits separated by mutational regime. We used the ESM-2 model size with the top PPO win rate from panel **a** (model size indicated in parentheses). The parent sequence’s predicted function value is depicted with the dashed line. **c**, Parent protein structures predicted by AlphaFold3^40^ overlaid with mutational frequency of sequence designs from PPO ESM-2 relative to pre-trained ESM-2 with interesting high confidence mutations labeled. **d**, Win rate of sequence designs from pre-trained, SFT, and PPO ESM-2 (650M) relative to pre-trained ESM-2 (650M) in low-N settings where reward models were trained with up to 1000 sequence variants.

The largest gains often came from the SFT phase, with additional but more modest improvements from PPO. In some cases, PPO offered little gain because the SFT model had already reached near-perfect win rates, while in others, this plateauing may reflect our limited hyperparameter optimization (Supplementary Table 6-7). Analyzing the reward scores of sequences generated at each training stage, we found that RLXF-aligned models consistently sampled sequences with higher predicted function than both the wild-type and those generated by the base ESM-2 (Fig. 2b). These models also sampled sequences outside the distribution of the reward model training data, exhibiting both more mutations and higher predicted function values (Fig. 2b). Notably, RLXF enabled the generation of high-reward variants extrapolating up to 20 mutations, while still outperforming the base model (Supplementary Figure 9).

To understand how RLXF integrates PLMs’ evolutionary priors with functional signals from reward models, we analyzed how the model’s amino acid preferences changed throughout the RLXF training process. Specifically, we examined the distribution of single-mutant logits for wild-type sequences across each alignment stage (Supplementary Figure 10–11). For proteins with broad evolutionary coverage (i.e. from large, diverse sequence families), RLXF increased positional entropy and decreased the maximum predicted amino acid probability at each site. This suggests the base PLM encoded strong evolutionary biases, which RLXF shifted to explore novel, functionally driven mutations. In contrast, for proteins with shallow evolutionary coverage, RLXF reduced entropy and increased confidence, suggesting the model became more focused by learning new functional constraints from the reward model. Bgl3, despite belonging to a large and well-represented protein family, showed decreased entropy and increased confidence during RLXF training. This likely reflects strong alignment between its reward model training data and evolutionary patterns^36^, causing RLXF to reinforce known substitutions rather than explore novel ones.

The base ESM-2 and the RLXF-aligned models exhibit distinct mutational preferences, reflecting a shift from evolutionarily conserved substitutions to mutations that enhance experimentally measured function (Fig. 2c). In GB1, which binds mammalian IgG, the RLXF-aligned model strongly favored substituting V29 with a positively charged residue near the binding interface. Structural analysis of the GB1–IgG complex suggests this substitution enables formation of a salt bridge with a nearby aspartate on IgG. Because this acidic residue is not conserved across IgG homologs, evolution-based PLMs do not favor this salt-bridge interaction. In Ube4b, RLXF favored mutations D68N, D139N, and N142T that are known to enhance auto-ubiquitination or stabilize the active E2∼Ub–E4B complex. Although these mutations may drive oncogenic activity and are disfavored by evolution, RLXF selected them for their biochemical benefit, decoupling functional optimization from evolutionary constraint. In Pab1, RLXF prioritized the Q94C mutation, the top enriched mutation in the reward model training data. However, the original DMS study identified Q94C as a false positive, with no measurable impact on growth. This illustrates how experimental artifacts in the training data can mislead the model during alignment, influencing sequence generation in unintended ways. In avGFP, RLXF enriched for well-characterized mutations (V163G, D180E, and T108G) that are known to enhance brightness and structural stability. In Bgl3, it prioritized A150D, a thermostabilizing substitution previously shown to boost enzyme performance under heat stress. Together, these results show that RLXF systematically shifts mutation preferences away from evolutionarily conserved residues and toward functional improvements (Supplementary Figure 12). This enables the generation of sequences with enhanced properties beyond what is accessible through natural variation.

Most protein engineering campaigns operate with limited experimental data. We therefore evaluated whether RLXF can guide functional sequence generation using reward models trained on reduced numbers of characterized variants. For the reward model, we used simple linear regression augmented with evolutionary features (Supplementary Figure 13-19; Table 10), which performs well in low-data settings^29^. Functional alignment with as few as 10-100 variants reliably shifted generation toward higher predicted fitness and improved win rates over the base ESM-2 (650M) model (Fig. 2d)

Protein families with greater natural diversity consistently showed stronger gains from RLXF with low-N reward models. This trend was supported by a direct correlation between a family’s number of effective sequences (*N*_*eff*_) in the UniRef100 database and win-rate improvement across the six protein families analyzed (Supplementary Figure 20). While the reward model does contain evolutionary features trained on family MSAs, we found little correspondence of reward model performance with *N*_*eff*_. Instead, we hypothesize the effect comes from differences in the calibration, strength, and stability of ESM-2’s evolutionary prior.

We evaluated the base ESM-2 model’s evolutionary prior using three metrics: wild-type retention under masking to assess logit calibration, entropy inflation to assess confidence stability, and the self-consistency gap to assess prediction consistency (Supplementary Figure 21-25). GB1 and avGFP, which have limited evolutionary representation, showed poor calibration and stability across all metrics including low wild-type probabilities, high uncertainty upon masking, and inconsistent predictions. These results suggest that RLXF alignment is less effective for proteins with weak or unstable priors, even when reward models are accurate, because the model lacks a reliable foundation for functional steering.

### Generative design of CreiLOV fluorescent proteins

CreiLOV is an oxygen-independent fluorescent reporter protein, uniquely suited for use in hypoxic or anaerobic environments where the ubiquitous, oxygen-dependent green fluorescent protein (GFP) is non-functional^41,42^. This makes CreiLOV a promising tool for applications in low-oxygen settings such as gut microbiomes, tumor microenvironments, and high-density fermentations^43^. However, natural and engineered variants of CreiLOV remain substantially dimmer than GFP limiting their practical utility^34,41,44^. Importantly, CreiLOV and other LOV domain-containing proteins are blue-light photoreceptors in algae, fungi, and plants, and fluorescence is not part of their biological role. As a result, the natural sequence record and pLMs trained on it are unlikely to encode information needed to enhance this non-natural, yet biotechnologically useful, fluorescence property. Designing brighter CreiLOV variants therefore requires explicitly guiding models toward functional objectives beyond natural evolution.

We found pre-trained ESM models favor the natural CreiLOV sequence over mutations known to enhance fluorescence, highlighting their bias toward evolutionary conservation rather than functional optimization. For instance, the C43A mutation in the FMN-binding pocket is a rationally engineered substitution that disrupts the native photocycle and significantly increases fluorescence emission^44^. ESM models consistently preferred the native cysteine at position 43, favoring restoration of the original photoreceptor function at the expense of fluorescence (Supplementary Figure 26-28). Similarly, most pre-trained models did not favor the T7S mutation, the brightest single variant identified in the CreiLOV DMS dataset^34^. An exception was the ESM-2 (650M) model, which showed a modest preference for T7S, possibly reflecting model size effects or stochastic differences in pre-training that incidentally aligned with fluorescence-enhancing features (Supplementary Figure 27). These results underscore the limitations of relying on pre-trained models alone for non-natural function optimization.

We applied RLXF to CreiLOV using a reward model trained on CreiLOV DMS data to predict the fluorescence of variant sequences. Among the pre-trained models tested, ESM-2 (650M) showed the highest baseline performance in terms of both mean and maximum predicted fluorescence and was selected as the base model for alignment (Supplementary Figure 2). We also evaluated a variational autoencoder (VAE) trained on a multiple sequence alignment of natural CreiLOV homologs, given the previous success of a VAE generating diverse and functional luciferase variants^30^. ESM-2 models were aligned using our two-stage RLXF procedure: SFT followed by PPO. We skipped supervised fine-tuning the VAE because our VAE already generated sequence variants similar in sequence identity to CreiLOV, suggesting the VAE did not require SFT to initialize parameters for alignment (Supplementary Figure 29).

Functional alignment with RLXF significantly improved the sequence generation capabilities of both models (Fig. 3a). The aligned ESM-2 model achieved a 100% win rate over its base counterpart and generated sequences with the highest maximum, mean, median, and minimum predicted fluorescence (Supplementary Table 11). Moreover, RLXF alignment enabled ESM-2 to generate high-reward sequences containing up to 45 mutations, substantially diverging from the wildtype while maintaining enhanced predicted function (Fig. 3b). These results demonstrate the effectiveness of RLXF in driving functional innovation beyond what pre-trained models or evolutionary priors alone can achieve.

**Figure 3.**
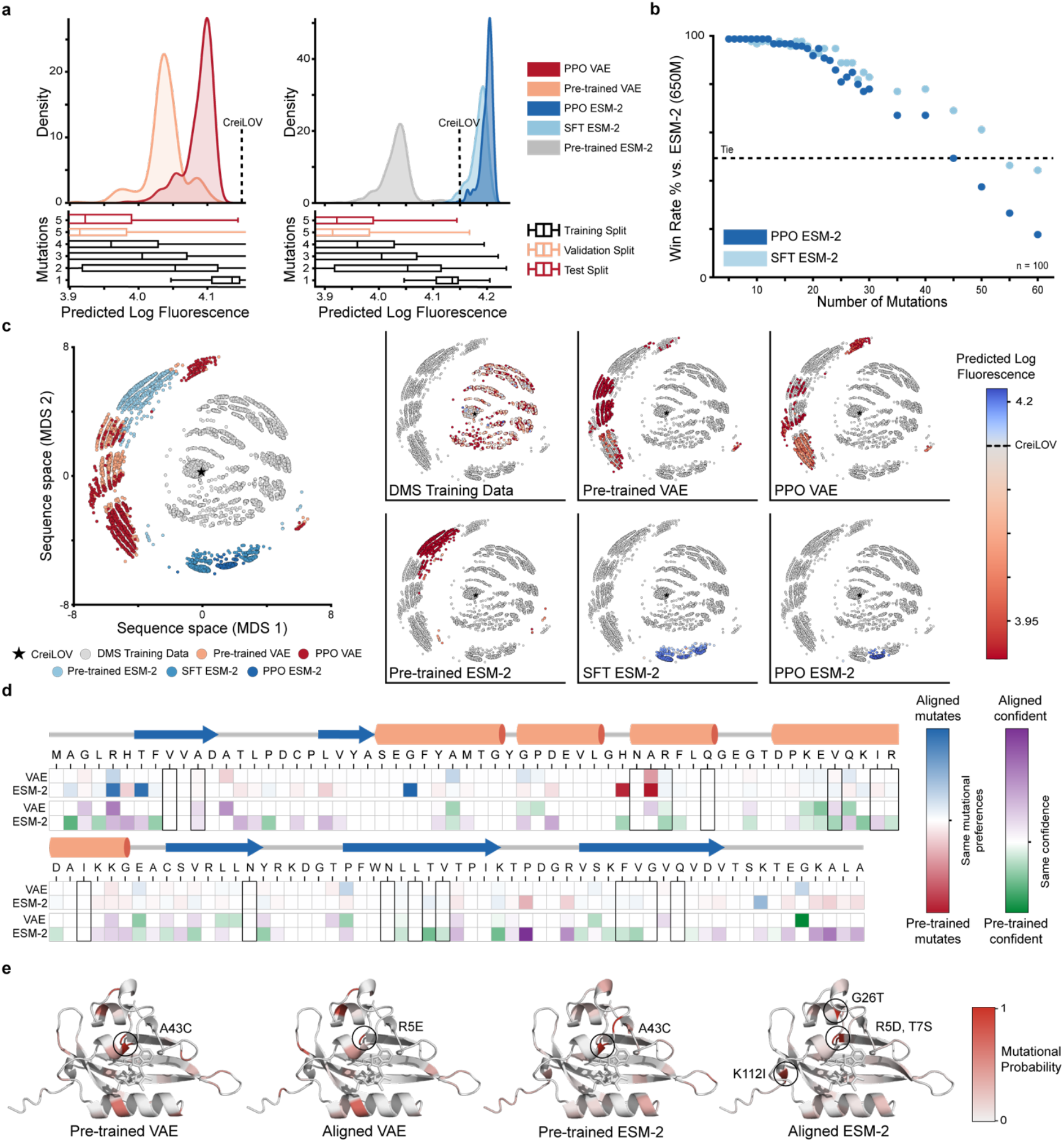
Aligned models learn to generate more fluorescent CreiLOV variants. **a**, Kernel density estimate plots depicting the distribution of predicted log fluorescence for 1,000 5-mutant designs generated by VAE or ESM-2 (650M) at each stage of alignment and boxplots for predicted function of sequences in reward model training, validation, and test splits separated by mutational regime. The CreiLOV parent’s predicted log fluorescence is depicted with the dashed line. **b**, Win rate for sequence designs with varying number of mutations generated from SFT or PPO ESM-2 (650M) relative to pre-trained ESM-2 (650M). **c**, Multi-dimensional scaling (MDS) visualization of the sequence from DMS data used to train the reward model and sequences sampled by the generative models. The generative models extrapolate beyond the DMS data, and the aligned models sample high fluorescence regions of the landscape. **d**, (top rows) Mutational frequencies across 30 sequence designs from the pre-trained and aligned models show how likely each model is to introduce mutations at specific positions in the CreiLOV sequence. (bottom rows) Shannon entropy calculated across 30 sequence designs from the pre-trained and aligned models reflects the diversity of amino acids selected at each position in the CreiLOV sequence. **e**, Mutational probability for pre-trained and aligned models mapped on the CreiLOV protein structure predicted by AlphaFold3^40^.

We observed distinct differences in the sequences generated by each model, indicating that they prioritize different regions of sequence space (Fig. 3c, Supplementary Figure 12). The pre-trained VAE and ESM-2 models showed some overlap in their sampling behavior, likely because both were trained on natural sequences. In contrast, the RLXF alignment process, particularly the SFT phase, dramatically shifted the sampling distribution of ESM-2 toward regions associated with high fluorescence. The PPO phase further refined this focus, concentrating sampling within these high-reward regions. To better understand how each model explores sequence space, we analyzed amino acid diversity across generated sequences at each position. The aligned ESM-2 model consistently favored the non-natural C43A substitution and introduced a combination of the non-natural mutations T7S, R5D, G26T, and K112I that were absent or rare in pre-trained models (Fig. 3d, Supplementary Figure 26-29). Shannon entropy differences further revealed that aligned ESM-2 made lower-entropy, and therefore more confident, predictions for these mutations strongly associated with enhanced fluorescence and did not disrupt critical flavin-binding residues except C43A (Fig. 3d). Mapping mutational probabilities onto the CreiLOV structure revealed that pre-trained models retained low mutational probabilities across many positions, whereas aligned models specifically increased mutational probabilities at key sites, highlighting a more targeted and confident exploration of beneficial sequence space after RLXF alignment (Fig. 3e, Supplementary Figure 30).

### Functionally aligned models generate high-performing CreiLOV variants

We characterized CreiLOV sequences generated from pre-trained and functionally aligned models to evaluate whether functional alignment improves sequence generation and enables discovery of variants with properties beyond those observed in nature. We selected a total of 60 designs: ten 5-mutant variants from the pre-trained VAE, ten 5-mutants from the aligned VAE, ten 5-mutants from the pre-trained ESM-2, ten 5-mutants from the aligned ESM-2, ten 10-mutants from the pre-trained ESM-2, and ten 10-mutants from the aligned ESM-2. We also included CreiLOV and the brightest known variant CreiLOV-T7S^34^ as controls. We expressed variants in *E. coli* BL21(DE3) and measured total cellular fluorescence to assess functional performance.

We found RLXF-aligned models consistently generated sequences with enhanced fluorescence compared to their pre-trained counterparts (Fig. 4a, Supplementary Figure 31). Both 5-mutant and 10-mutant designs from aligned ESM-2 showed statistically significant increases in mean fluorescence across the ten sampled variants relative to pre-trained ESM-2. The distribution of fluorescence values across all designs was bimodal with a clear separation point around 69% of CreiLOV’s fluorescence (Supplementary Figure 32). Based on this, we defined three fluorescence categories: low (<69%), medium 69–100%), and high (>100%) relative to CreiLOV. Over 80% of sequences generated by the pre-trained models fell into the low-fluorescence category, while the aligned models predominantly sampled sequences in the medium and high categories (Fig. 4b). Notably, 6/10 5-mutants from the aligned ESM-2 model exceeded CreiLOV fluorescence and the remaining 4 fell into the medium category. The aligned ESM-2 generated three sequences with fluorescence values greater than CreiLOV-T7S, the brightest known flavin-binding fluorescent protein (Fig. 4a). We also found that rationally combining the top 5 or 10 beneficial single mutations^34^ produced non-functional variants (Supplementary Figure 33). This highlights RLXF’s ability to learn synergistic, epistatic mutation combinations that cannot be identified by simply adding the effects of individual mutations.

**Figure 4.**
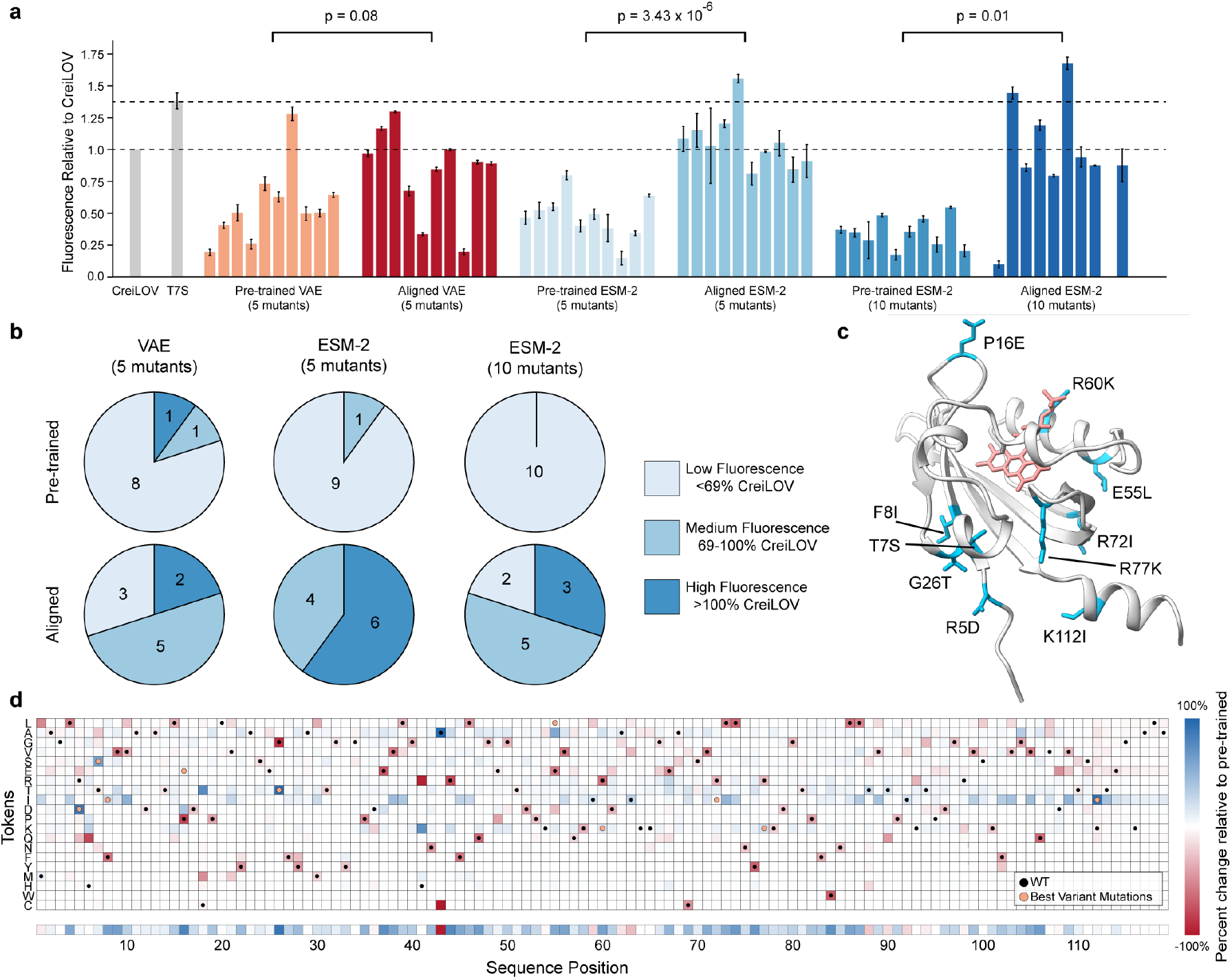
Experimental characterization of pre-trained and RLXF-aligned CreiLOV variants. **a**, Bar plot showing fluorescence relative to CreiLOV for variants generated by pre-trained and aligned ESM-2 and VAE models with 5 or 10 mutations per design. Error bars represent standard deviation (n=3). CreiLOV and T7S are shown in grey. Dashed lines indicate CreiLOV and CreiLOV-T7S average fluorescence values. Statistical significance determined by a two-tailed t-test between pre-trained and aligned model conditions is shown. **b**, Proportion of variants categorized as low, medium, and high fluorescence. **c**, AlphaFold3^40^ structure of the most fluorescent 10-mutant, with all 10 mutations highlighted in cyan and the FMN cofactor displayed in coral. **d**, Comparison of amino acid probabilities between RLXF-aligned and pre-trained ESM-2 (650M) models, highlighting mutational preferences learned during RLXF alignment. The bottom row shows the difference in cumulative probabilities assigned to non-CreiLOV amino acids between the aligned and pre-trained models, highlighting how likely each model is to introduce mutations at each position.

Our top three RLXF-generated variants exhibited higher total cellular fluorescence than CreiLOV-T7S. Total cellular fluorescence depends on several factors, including protein expression level, FMN binding affinity and kinetics, and intrinsic optical properties such as extinction coefficient and quantum yield. We therefore sought to determine which of these factors contributed to the improved fluorescence of the RLXF variants. Measurements of intrinsic photophysics showed that the RLXF variants have quantum yields of ∼0.6, which is higher than wild-type CreiLOV but similar to CreiLOV-T7S. Because their quantum yields are comparable to CreiLOV-T7S, the additional gains in cellular fluorescence are likely explained by other factors, such as improved FMN binding, more efficient chromophore loading or maturation, or increased protein expression.

RLXF largely preserves the mutational landscape learned during ESM-2 pretraining while sharpening amino-acid preferences at fluorescence-relevant positions (Fig. 4c-d). For example, the aligned model shifts position 43 from the evolution-favored cysteine to alanine, a rationally designed substitution known to improve CreiLOV fluorescence. RLXF also consistently prioritized a core set of high-confidence mutations (R5D, T7S, G26T, K112I). The brightest RLXF design contains 10 mutations and shows a 1.2-fold increase in fluorescence relative to the previously best variant, CreiLOV-T7S (Fig. 4c). This variant combines the four high-confidence substitutions with additional mutations that were ranked very low or absent in the DMS dataset. Specifically, F8I, P16E, E55L, R60K, and R72I ranked 1376th, 105th, 330th, 53rd, and 986th, respectively, and R77K was not present in the DMS data. These mutations would be difficult to prioritize using single-mutation metrics alone. Together, these results show that RLXF leverages evolutionary pretraining for sequence diversity while using experimental feedback to discover synergistic, non-obvious combinations of mutations that improve fluorescence.

## Discussion

In this work, we introduced Reinforcement Learning from eXperimental Feedback (RLXF), a general framework for aligning protein language models with human-defined functional objectives. Drawing inspiration from advances in aligning large language models with human preferences, RLXF uses a reward model based on experimental data to guide sequence generation toward desired biophysical or biochemical properties. We demonstrated the effectiveness of RLXF by applying it to five diverse protein systems, showing that functional alignment consistently improves the generation of high-performing sequences beyond what pre-trained models can achieve. We used the oxygen-independent fluorescent protein CreiLOV to experimentally demonstrate aligned models generate more functional sequences with greater overall fluorescence values than pre-trained models. Together, these results show that functional alignment can substantially expand the design capabilities of pLMs beyond the constraints of natural evolution.

The reward model is central to aligning generative pLMs with human protein engineering objectives. It provides feedback to the generative model, biasing its sampling toward sequences with desirable properties. In principle, the reward model in RLXF can be any sequence- or structure-based scoring function, including zero-shot predictors^7,45^, evolutionary Potts models^46,47^, Rosetta-based energy functions^48,49^, solubility predictors^50^, or immunogenicity predictors^51,52^, or even multi-objective criteria. In this work, we use supervised models trained on experimental sequence-function data because they offer highly accurate predictions of complex protein properties^27,28^. Importantly, by learning directly from experimental data, these models guide sequence generation toward the desired functional outcomes, even when the underlying biophysical mechanisms are poorly understood.

RLXF takes a general-purpose pLM, originally trained on evolutionary sequence data, and endows it with an explicit notion of biochemical function. We find RLXF shifts the model’s mutational preferences away from simply mimicking evolutionarily related sequences and towards prioritizing mutations supported by experimental functional data. Our analysis of mutations preferred by RLXF-aligned models shows alignment drives generation to optimize sequences according to the reward model data, balancing evolutionary constraints with experimentally defined functional objectives. RLXF achieves substantial win-rate gains with fewer than 100 labeled sequence variants for the reward model when the pre-trained pLM prior is well calibrated. Proteins with weak priors require improved calibration in this low-N setting. Calibration can be improved using biophysical simulation data such as Rosetta-derived simulation datasets previously shown to significantly improve the predictive performance of finetuned ESM-2 for avGFP and GB1 sequence variants in low-N settings^11^. Together, these results emphasize alignment quality depends on the reward signal and the prior: high-quality experimental data matters because you get what you optimize for and well-calibrated evolutionary priors are critical for reliably steering generation toward functional enhancement.

The two phases of RLXF training, supervised fine-tuning (SFT) and proximal policy optimization (PPO), work together to reshape the pLM through a combination of grounding and controlled exploration. During SFT, the model learns mutational patterns that are strongly associated with improved function, anchoring it in a favorable region of sequence space. PPO then further refines the model by encouraging the discovery of additional beneficial mutations and by reducing overfitting to the SFT training set through broader sampling and evaluation. This exploration step is critical because supervised fine-tuning alone has been shown to limit generative diversity and fail at out-of-distribution generation tasks^53^, making PPO essential for producing a robust, functionally aligned generative model.

The RLXF-aligned ESM-2 models generated multiple sequences with greater fluorescence than the CreiLOV parent. We hypothesize the best-performing 10-mutant achieves its 1.7-fold fluorescence enhancement through a synergistic combination of mutations that optimize placement of the FMN cofactor while rigidifying the overall protein structure in a fluorescence-competent conformation. Four core mutations (R5D, T7S, G26T, K112I), consistently observed across high-performing variants, likely stabilize the chromophore environment, consistent with previous findings that identified T7S as the brightest CreiLOV variant^34^. However, our variant exceeds CreiLOV-T7S, likely due to additional mutations (P16E, F8I, E55L, R60K, R72I, R77K) that individually rank lower in the DMS data but act cooperatively to further enhance cellular fluorescence^34^. Structural analysis indicates that these mutations do not substantially reposition FMN but instead reduce pocket solvation and slightly rewire chromophore-protein hydrogen bonding. Notably, the R60K mutation selectively weakens a guanidinium–FMN interaction while preserving overall pocket packing. These strategically positioned mutations likely rigidify the chromophore pocket, creating a more hydrophobic environment that reduces quenching collisions, a mechanism consistent with previous observations linking chromophore rigidification to improved quantum yield. This highlights how RLXF can uncover combinatorial effects that are difficult to predict from single-mutant data alone.

Direct Preference Optimization (DPO) is a simpler alternative to reinforcement learning from human feedback (RLHF), eliminating the need for a separate reward model and policy optimization^54^. Instead of using reinforcement learning, DPO directly fine-tunes the model to prefer outputs ranked higher by human feedback. However, RLHF consistently outperforms DPO for complex generation tasks in NLP because it allows the model to explore and optimize beyond the fixed preference dataset^55,56^. With RLHF and policy optimization methods like PPO, the training distribution can shift dynamically as the model samples new outputs, receives feedback, and adapts, enabling discovery of higher-quality solutions not present in the original data. This ability to actively explore and extrapolate is critical when optimizing over complex, poorly mapped spaces. As a result, PPO-based RLHF remains the method of choice for training top foundation models, despite its greater computational cost and lower training stability compared to DPO.

Several recent studies have applied DPO to align pLMs with functional objectives^1,20,57–59^. Hie et al. introduced ProteinDPO, applying DPO to a structure-conditioned pLM (ESM-IF1^60^) to align sequence generation with experimental measurements of thermostability^57^. They demonstrated that DPO alignment improved stability prediction across diverse datasets, generalized to new tasks like binding affinity prediction, and enabled *in silico* generation of more stable protein sequences, although they did not experimentally validate newly designed proteins. Stocco et al. developed DPO_pLM, adapting DPO for autoregressive pLMs and demonstrating its effectiveness in optimizing properties such as structure quality and binding affinity^58^. They experimentally validated their approach by designing epidermal growth factor receptor (EGFR) binders, several of which achieved nanomolar affinity, demonstrating functional enhancement over the wildtype ligand. Together, these studies show that DPO-based fine-tuning can improve the functional capabilities of pLMs, although exploration is limited to the regions defined by the available preference data, in contrast to reinforcement learning approaches like PPO that allow active discovery beyond the training set^53,55,56^.

RLXF is a general-purpose framework that can be applied to proteins with diverse folds and functions. It is accessible to new users and can be implemented with modest computational resources. Our typical workflow takes approximately three days: one day to train a reward model, one day to generate a high-reward sequence dataset for SFT via simulated annealing, and one day to perform SFT and PPO on a single GPU. We recommend training a simple sequence-to-function reward model. In our work, we used an ensemble of fully connected neural networks that take one-hot encoded amino acid sequences as input and output predicted functional scores. We also recommend testing multiple open-source pre-trained pLMs of varying sizes to identify the base model most amenable to functional alignment. Across our experiments, the ESM-2 (650M) model generally achieved the best overall performance, although the optimal model size varied depending on the specific protein system. Regardless of model size, we strongly recommend using parameter-efficient fine-tuning methods, as they consistently outperformed full-parameter fine-tuning by reducing catastrophic forgetting of the pre-trained model’s knowledge during both SFT and PPO.

The growing success of functionally aligned pLMs signals a broader shift from passive extraction of information encoded in natural sequences to active, experimental steering of generative models toward human-defined engineering goals^1,2,20,57,58,61–63^. RLXF offers a general and flexible framework to align pLMs with virtually any measurable property, enabling the rapid generation of diverse high-fitness sequence libraries. Looking ahead, RLXF and related approaches could be integrated into iterative, multi-objective workflows, where models are continuously refined through experimental feedback in autonomous self-driving laboratories^64,65^. By combining the power of pre-trained models with experimental feedback, we can open vast regions of sequence space and design proteins with entirely new functions beyond anything produced by natural evolution.

## Methods

### Data curation for reward models

We used an extensive CreiLOV DMS dataset^34^ to train reward models. The training split contained 2,204 single mutants (92.6% coverage), 176 double mutants, 978 triple mutants, and 3,565 four mutation variants. The validation and test sets were curated using a 75/25 data split for the 9,603 five mutation variants. We used similar data splits for reward models trained to guide the alignment of ESM-2 across diverse protein classes (Supplementary Table 2).

### Reward model training

We trained 100 multi-layer perceptrons with fixed training, validation, and test data splits to predict the log mean fluorescence of CreiLOV sequence variants in a supervised manner using a mean squared error (MSE) loss objective. Each multi-layer perceptron in the ensemble was initialized with a different seed. Input sequences were one-hot encoded. Hyperparameters were the same for each multi-layer perceptron reward model (Supplementary Table 3).

### Data curation for VAE

We curated 260,349 natural sequences related to CreiLOV from the protein database UniRef90^66^ with the Hidden-Markov model homology search tool Jackhmmer^67^ to obtain a multiple-sequence alignment (MSA) containing proteins related to CreiLOV with a maximum of 2 iterations (N=2). We removed sequences less than 75% of the length of CreiLOV^45^, removed sequences with an amino acid repeating 10 times in a row, and removed positions of the MSA not corresponding to CreiLOV^30^. We reweighted the remaining 243,682 sequences with neighbors classified as having a Hamming distance/length of sequence greater than 0.98 to reduce phylogenetic bias from uneven sampling^45,68,69^. We withheld 100 sequences from the reweighted MSA to later assess VAE overfitting to the training set as a pseudo-test set given that these sequences are unlabeled. We randomly sampled the remaining 243,582 sequences with a 90/10 split for the training and validation sets.

### VAE pre-training

The VAE is trained by maximizing a modified version of the Evidence Lower Bound (ELBO) that effectively minimizes the Kullback–Leibler divergence between the variational approximation and the true posterior distribution^45^ as shown with equation (1):

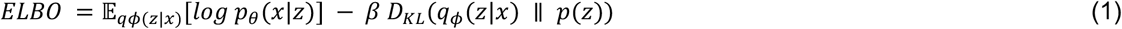

The first term can be considered a reconstruction loss that is computed using cross-entropy between an input one hot-encoded sequence and output likelihoods. β is the weight for the Kullback–Leibler divergence term *D*_*KL*_ with a prior distribution *p*(*z*) of *N*(0, *I*). Hyperparameters were optimized via a grid search of 673 combinations (Supplementary Table 8).

### Generating supervised fine-tuning sequence datasets

We generated sequence datasets for supervised fine-tuning via 100 simulating annealing simulations by adapting code from literature^28^3/6/26 3:51:00 PM. We designed variants of a fixed number of mutations and high predicted function according to our reward models. At each step of simulated annealing, we introduced 2 mutations at a time sampled from a Poisson distribution (*λ* = 1) with the number of mutations constrained. We predicted the function of variants using a conservative, 5th percentile, score from the ensemble of reward models. Mutations that enhanced function were automatically accepted while all other mutations were accepted with a probably calculated with equation (2):

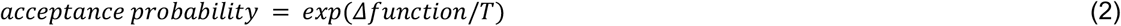

where *T* decreases with a logarithmic temperature gradient selected to have an initial acceptance probability near 40% for mutations that do not increase function to encourage early exploration and approaches 0% towards the end of 50,000 steps (Supplementary Table 5). We removed duplicate sequences from the final sequences from trials to obtain a sequence dataset for supervised fine-tuning.

### Supervised fine-tuning ESM-2

We supervised fine-tuned ESM-2 using the standard masked language modeling (MLM) objective. We masked positions of each sequence in the sequence dataset with mutations relative to the natural CreiLOV sequence prior to the mutation C43A^44^. During training, we compute the cross-entropy loss for masked residues, minimizing equation (3):

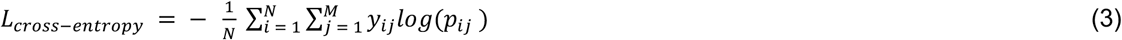

where *N* is the total number of masked positions in a mini-batch, *M* is the size of the amino acid vocabulary for the ESM-2 tokenizer, *y* is the one-hot indicator for the true amino acid at the masked position *i* (token *j*), and *p* is the ESM-2’s predicted probability for that amino acid.

We trained parameters in the language model head, the last transformer block, and the LayerNorm weights and biases in the second to last transformer block of ESM-2 (650M), keeping the rest of the parameters frozen. We used the Adam optimizer with a base learning rate of 0.00511 and weight decay of 0.00351. We grouped parameters into language model head and transformer layer-specific parameter groups, initialized with the base learning rate and decayed by a factor of 0.89 for each successive group. We wrapped Adam in a CosineAnnealingWarmRestarts scheduler (*T*_0_ = 10, *T*_*mult*_ = 1) over a single-epoch run with a batch size of 8. We applied a gradient norm clipping with a max norm of 3. Hyperparameters were selected by Optuna^70^ after 1,000 trials of hyperparameter optimization with a Tree-Parzen Estimator (TPE) sampler and a custom PyTorch callback to terminate trials if model parameters became non-finite (i.e., NaN or infinity) after each min-batch (Supplementary Table 6).

### Generating sequence variants with top p-sampling thresholds

We generated sequence variants in 2 distinct steps, attempting to mimic how a directed evolution campaign proceeds. First, we masked CreiLOV one position at a time and generated the probabilities for amino acids at each position. If the probability of any non-natural amino acid exceeded our high confidence threshold, we introduced this mutation into all sequence variants. Then, we masked CreiLOV with the high confidence mutations one position at a time and generated the probabilities for amino acids at each position. We computed the sum of probabilities for non-natural mutations at each position. We sampled positions weighted by the cumulative probability of non-natural mutations. We used a second cumulative probability threshold to avoid sampling positions where mutations were unlikely to enhance function. For a sampled position, we sampled amino acids weighted by the probability of each amino acid including the natural amino acid. We iteratively sampled positions until the desired number of mutations was achieved.

### Proximal policy optimization

Our PPO implementation utilizes two copies of SFT ESM-2 or the pre-trained VAE weights. One copy remains frozen while the other model has trainable parameters. In agreement with reinforcement learning terminology, we refer to the trainable generative model as a policy in this section. Proximal policy optimization relies on the minimization of the clipped surrogate objective function in equation (4):

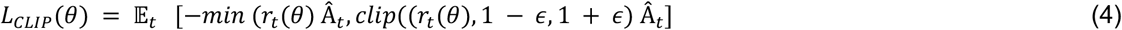

where the probability ratio *r*_*t*_(*θ*) is computed between the trainable policy π_θ_ and the frozen model π_θ,*ref*_ as shown in equation (5):

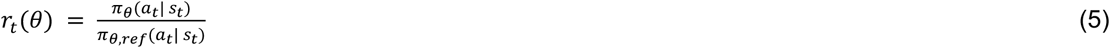

where the ratio provides a comparison of probabilities from the frozen and policy models for the amino acid mutations *a*_*t*_sampled by the policy model at state *s*_*t*_. The hyperparameter ϵ in equation (4) establishes a pessimistic bound on the probability ratio to prevent excessively large model updates and to help balance exploration and exploitation. Our total reward *R*(*x, y*) is computed with equation (6):

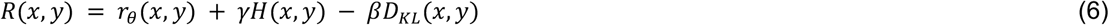

where the parent sequence prompt to the policy is *x* and the sequence variants with sampled amino acid mutations are *y*.

We normalize the predicted functional reward *r*_*θ*_(*x, y*) in a manner specific to our task of learning to generate protein sequence variants with function-enhancing mutations relative to a parent sequence. Similar to GRPO^71,72^, we do not rely on a critic model^73^ to estimate the advantage Â_*t*_. We instead estimate the advantage Â_*t*_ by generating multiple sequence variants *y* for the same parent sequence *x* and use the parent sequence as a group baseline to reduce the variance of predicted rewards used for policy gradients. We first incorporate policy knowledge with the same sampling strategy used when generating sequence variants with top p-sampling thresholds to focus on exploring mutations more likely to enhance function. We compute a conservative predicted function reward for each sequence variant using the 5th percentile score from the ensemble of reward models and take the max of conservative predicted function rewards for the batch to provide an estimate of the state of our policy for generating sequence variants with enhanced function. We finally divide this normalized reward by the predicted function of the parent sequence prompt. We found these simplifications allowed us to estimate Â_*t*_ with *R*(*x, y*) and increase the predicted function of sequence variants after PPO. We calculate the average pairwise Hamming distance term *H*(*y*) between generated sequence variants for each batch to encourage the generation of diverse sequences with equation (7):

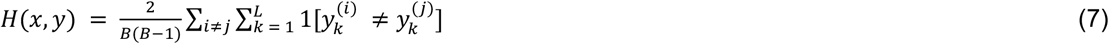

where *B* is the number of generated sequence variants in the batch, *L* is the length of the variant sequences, amd 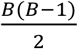 is the number of unique pairings. We weight the average pairwise hamming distance term with *γ*. We calculate the Kullback–Leibler divergence penalty *D*_*KL*_ between the policy and frozen model to prevent the policy from drastic changes relative to the frozen model with equation (8):

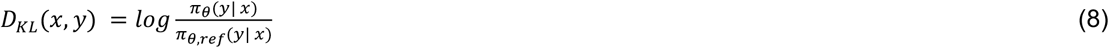

where *y* is generated sequence variants for the parent sequence prompt *x*. When aligning ESM-2, we iteratively masked each parent sequence position to generate log probabilities for single amino acids mutations from the frozen model prior to training. This time-consuming calculation requires a forward pass for each position of the parent sequence prompt but does not need to be repeated for the frozen model. We calculated new log probabilities each epoch of PPO for the trainable policy. We used this method to simplify and expedite the calculation of *D*_*KL*_ each epoch. We weight this term with *β*.

Our variant of PPO for the alignment of the masked language model objective pLM ESM-2 can be summarized with the following steps:

1. Iteratively mask positions of the parent sequence and generate amino acid log probabilities with the frozen policy for amino acids at each position
2. Iteratively mask positions of the parent sequence and generate amino acid log probabilities with the trainable policy for amino acids at each position
3. Calculate *D*_*kL*_ between log probabilities from steps 1-2
4. Mask all positions of the parent sequence where an amino acid mutation exists with a probability greater than a high confidence threshold according to the trainable policy and generate amino acid probabilities for those positions from the policy and frozen models.
5. Add high confidence amino acids to parent sequence
6. Iteratively mask positions of the variant sequence with high confidence mutations and generate amino acid log probabilities with the trainable policy for amino acids at each position
7. Compute cumulative probability of amino acids mutations for each position
8. For each sequence in a batch, iteratively sample positions using cumulative probability weights and generate log probabilities with policy and frozen models. Sample amino acids weighted by the probability of each amino acid for the sampled positions according to the policy model. Repeat until the specified number of mutations are introduced
9. Compute the ratio *r*_*t*_ (*θ*) for amino acid mutations present in sequence variants with log probabilities from steps 4 and 8
10. Compute the total reward *R*(*x, y*) from sequence variants
11. Compute *L*_*CLIP*_(*θ*)
12. Minimize *L*_*CLIP*_ (*θ*) via gradient descent on trainable policy parameters
13. If the number of iterations were greater than 1 for an epoch, repeat steps 4, 8, 9, 11, and 12 to minimize *L*_*CLIP*_ (*θ*) via gradient descent on trainable policy parameters with updated *r*_*t*_(*θ*).
14. Repeat steps 2-13 for each epoch of PPO.

When aligning a VAE via PPO, we recursively added Gaussian noise *N*(0, σ^G^) to the mean latent representation of the parent sequence to identify the variance required to generate a batch of sequence variants with the desired number of mutations at the beginning of each epoch. We consider the likelihood matrices for a mini-batch of generated sequence variants from the trainable and frozen VAE policies to calculate *D*_*KL*_. The likelihood matrices are size 21 × *L* where 21 refers to the 20 amino acids and gap tokens possible for a sequence position and *L* refers to the length of the parent sequence prompt. We sample each column of a likelihood matrix from the trainable VAE policy to generate each sequence variant in a batch.

We implemented a parameter-efficient proximal policy optimization by training parameters in the language model head, the last transformer block, and the LayerNorm and dense output weights and biases in the second to last transformer block of ESM-2 (650M), keeping other ESM-2 parameters frozen to preserve ESM-2 pre-training knowledge important for out-of-distribution generation tasks^74^. Beyond the 1st epoch, we unlocked additional parameters to include parameters in the language model head and the last 5 transformer blocks in total. We adapted this strategy from a recent study fine-tuning ESM-2^11^. We used the Adam optimizer with a base learning rate of 0.00866 and weight decay of 0.00995. We grouped parameters into layer-specific groups, initialized with the base learning rate and decayed by a factor of 0.885 to assign a learning rate to each successive group. We wrapped Adam in a CosineAnnealingWarmRestarts scheduler (*T*_0_ = 1, *T*_*mult*_ = 2) over a 2 epoch run with an initial batch size of 1 that increased by 1 each epoch. We applied a gradient norm clipping with a max norm of 6.82 and smoothed parameter updates using an exponential moving averaging. We used the following hyperparameters for calculating *L*_*CLIP*_(*θ*): *ϵ* = 0.174, *γ* = 1.0e-06, *β*_1*st epoch*_= 1.0e-08, *β*_*beyond* 1 *st epoch*_ = 1.0e-07 (7,9).

### Curating multiple sequence alignments for EVE models

Following the EVE^75^ repository workflow (https://github.com/OATML-Markslab/EVE), we generated MSAs by iterative profile-HMM homology search using Jackhmmer against UniRef100 using the EVCouplings^76,77^ pipeline (https://github.com/debbiemarkslab/EVcouplings). We retained homologous sequences that aligned to at least 50% of the target sequence length and filtered alignment columns to require ≥70% residue occupancy. To select a final MSA per protein, we explored a range of thresholds and prioritized alignments with high target coverage and sufficient depth, favoring configurations with coverage *L*_cov_ ≥ 0.8*L* and total sequences *N* satisfying 10*L* ≤ *N* ≤ 100,000 consistent with the EVE guidance (Supplementary Table 10).

### Training EVE models

EVE models were trained for each protein following the EVE^75^ repository workflow (GitHub repository: https://github.com/OATML-Markslab/EVE).

### Obtaining Evolutionary Density Features from TranceptEVE

To provide this evolutionary feature consistently across protein families, we used TranceptEVE^78^ that achieves the best average Top-10% recall among sequence-based baselines on ProteinGym^79^. TranceptEVE combines outputs from the family-agnostic autoregressive pLM Tranception^80^ with a family-specific EVE model and leverages MSA retrieval at inference time. We curated MSAs from UniRef100 for each protein and trained family-specific EVE models that we integrated with Tranception to construct a TranceptEVE model per protein. We computed evolutionary density features using TranceptEVE as described in equation (9):

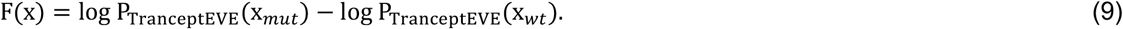

The log-likelihood at each position is obtained using the log likelihood log P_T_ (x_*i*_| x_<i_) from Tranception S^80^ (https://github.com/OATML-Markslab/Tranception), log P_EVE_(x_*i*_) from the protein family-specific EVE model we trained that is obtained by inputting into the VAE the wild-type sequence from which the MSA was acquired and then averaging the resulting log-softmax outputs from the decoder network across a large number of samples from the approximate posterior and from the distribution over decoder parameters as previously described, and log P_MAS_(x_*i*_) from the distribution over amino acids at each position of the retrieved MSA as described in equation (10):

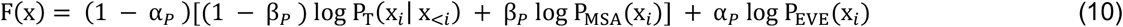

The constants α_*P*_ and β_*P*_ depend on MSA depth as previously recommended^78^.

### Data curation for augmented ridge regression reward models

We used the same train, validation, and test splits for reward models trained to guide the alignment of ESM-2 across diverse protein classes (Supplementary Table 2). We constructed training subsets by uniformly sampling the full training set at multiple target sizes while keeping the test sets fixed. For each target training size, we generated 10 independent replicates.

### Training augmented ridge regression reward models

We trained augmented ridge regression reward models to predict the experimental functional score from a combination of sequence identity features and an evolutionary density feature derived from TranceptEVE as previously described^78^. Each sequence variant was represented using per-position one-hot encoding of amino acid identities, and we appended a single continuous evolutionary feature^29^. Ridge regression minimizes the objective in equation (11):

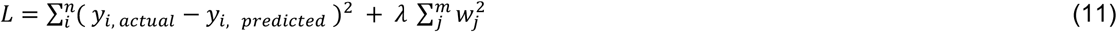

where *n* is the number of training samples, *m* is the number of features, and *w*_*j*_ are the model coefficients Models were implemented in scikit-learn with penalty strength of *λ* = 1. To tune the relative contribution of the evolutionary feature, we performed 5-fold cross-validation within each training subset when at least five training examples were available. We selected the evolutionary feature scaling/regularization setting from 10^−3^, 10^/-4^, 5 × 10^/-5^, 10^/-5^ by maximizing mean Spearman correlation across folds. We then retrained the final model on the full training subset using the selected setting.

### Low-N RLXF with augmented ridge regression reward models

For low-N functional alignment, we proceeded with 3 of 10 augmented ridge regression models trained with 10, 100, 200, 500, or 1000 combinatorial variants per protein. The only modification to our previously described alignment pipeline was the construction of the SFT training dataset. We extracted the top 40 highest-weighted non–wild-type amino acid substitutions from the trained ridge model (excluding stop codons), enumerated all combinatorial 5-mutation variants, and scored candidates with TranceptEVE to obtain the evolutionary density feature required by the ridge model. We then re-scored candidates using the same ridge model and selected a final set of 150 high-scoring variants. To reduce redundancy, we curated a diverse subset of 25 sequence variants using normalized Hamming-distance–based greedy farthest-point selection. These curated variants were used as the SFT training dataset. SFT optimization and all PPO procedures (policy parameterization, rollout strategy, reward computation, and hyperparameters) were otherwise identical to our main RLXF alignment protocol, with the augmented ridge regression model serving as the PPO reward function for the corresponding replicate.

### Computing shifts in pLM priors across SFT and PPO and calibration of pre-trained pLM priors

To quantify how RLXF changes *sequence priors*, we computed a set of calibration and distribution-shift metrics from ESM-2 logits at each stage of alignment. For each protein wildtype sequence *x* = (*x*_1_, …, *x*_*L*_), we computed masked-position logits using pseudo-log-likelihood scoring in equation (12) and full-sequence logits from a single forward pass without masking in equation (13).

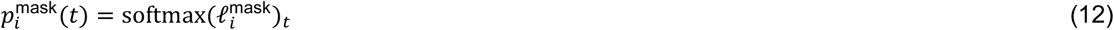

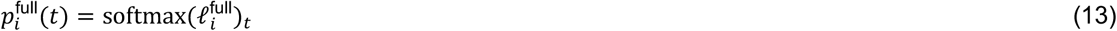

where 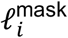 and 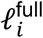 are the corresponding logits over the tokenizer vocabulary *V*at position *i*. Metrics were computed either on the wildtype sequence (*k* = 0) or along a high-confidence mutation trajectory 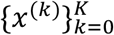, where *x*^(*k*)^contains the top-*k* substitutions whose probability exceeded a fixed probability threshold from the RLXF-aligned ESM-2. Below, 𝒜 denotes the 20 canonical amino acids.

For each stage (pretrained, SFT, aligned), we computed the metrics in equations (14-17) on the high-confidence mutation trajectory 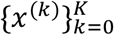 . This allowed us to quantify how local distributions and calibration evolve as increasingly many high-confidence substitutions are introduced, and to compare how SFT and PPO shift priors relative to the pretrained model under matched mutational regimes. To quantify how much the model’s local residue distribution shifts relative to a reference stage, we computed cosine similarity in equation (14) between AA-only masked distributions at each position and averaged across positions:

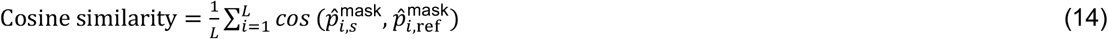

where 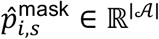is the masked distribution from equation (12) restricted to amino acids 𝒜 Values close to 1 indicate minimal distribution shift, while lower values indicate stronger drift away from the reference stage prior.

We measured local uncertainty under masked inference using the average positional entropy over amino acids in equation (15):

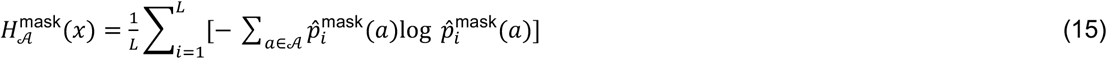

where 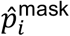 is derived from equation (12) restricted to amino acids 𝒜. Higher entropy indicates broader residue uncertainty, whereas lower entropy indicates sharper residue preferences.

To quantify distribution sharpness, we computed the mean maximum amino-acid probability per position in equation (16):

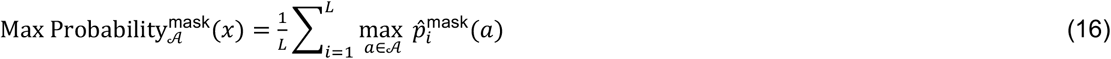

where 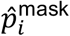 is derived from equation (12). Larger values indicate a more peaked distribution.

We computed pseudo-perplexity from the masked pseudo-log-likelihood of the observed residue at each position with equation (17):

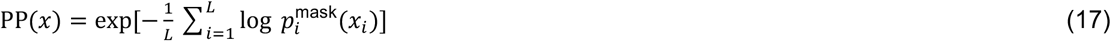

where 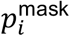 is defined in equation (12). Lower pseudo-perplexity corresponds to higher likelihood of the wildtype under masked inference and thus a stronger calibrated prior.

To quantify whether the wildtype residue is among the top-*K* predictions under masked inference, we computed equation (18)

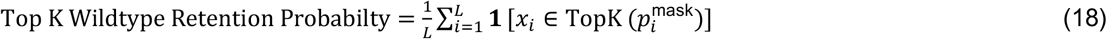

where TopK(*p*) returns the *K* most probable tokens under equation (12). We computed this metric for *K* ∈ {1, …, 20}. We also computed the wildtype rank gap as the change in the rank of the wildtype residue when comparing masked and full inference in equation (19):

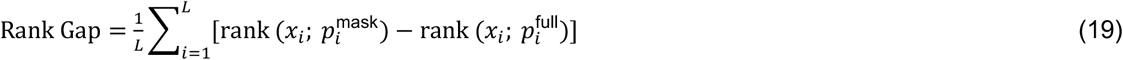

where rank(*x*_*i*_; *p*) is the 0-based rank of *x*_*i*_ when sorting tokens by probability under *p*(rank 0 is highest probability), and 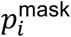 and 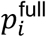 are defined in equations (12-13). Positive values indicate that masking decreases the WT rank on average.

We quantified sensitivity to masking using the self-consistency gap, defined as the difference in wildtype log-probability under full versus masked inference in equation (20):

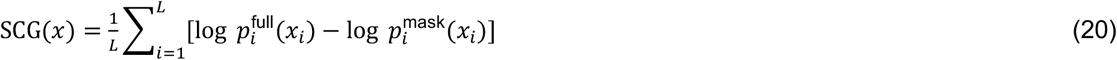

where 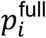 and 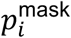 are defined in equations (12-13). Larger SCG values indicate stronger reliance on global context, or lack of local evolutionary understanding.

To measure how much uncertainty increases under masking, we computed entropy inflation as the difference between masked and full entropies over the full vocabulary *V* and took the difference in equation (21):

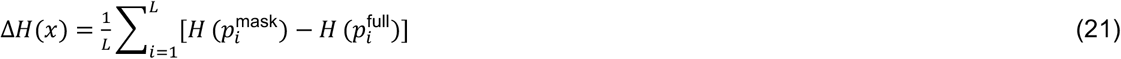

where the token-distribution entropy is in equation (22):

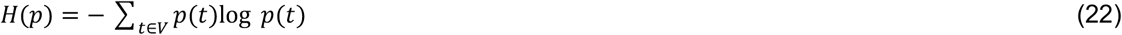

Here, 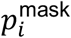 and 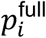 are defined equations (12-13). Larger Δ*H* indicates greater disruption of model confidence induced by masking.

### Direct preference optimization

In agreement with reinforcement learning terminology, we refer to the trainable generative model as a policy in this section. Direct preference optimization relies on the minimization of the objective function in equation (23):

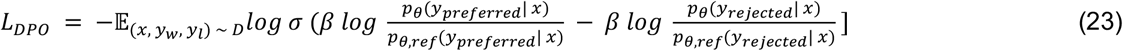

where *θ* is the policy with trainable parameters, *θ*_*ref*_ is the pre-trained model, *β* is a hyperparameter that determines the weight of the policy model relative to the pre-trained model, *x* refers to a mini-batch of sequences in a pre-processed sequence dataset *D, y* refers to the label of sequences in the mini-batch as preferred or rejected sequences based on their experimental function. *L*_*DPO*_ (*θ, θ*_*ref*_) compared the log-likelihood from the policy and pre-trained model and finetunes policy parameters to increase the ratio for preferred sequences relative to the ratio for rejected sequences. For the sequence dataset *D*, we split reward model training data into rejected and preferred sequences for each mutational regime using their experimental log fluorescence^34^. We pre-computed log-likelihoods for the entire sequence dataset for the pre-trained model to reduce training time as this remains constant throughout training. We implemented direct preference optimization with 2 sets of hyperparameters from literature^57,58^.

### Computing correlations

We computed the top 10% recall and Spearman rank correlation coefficient using the experimental log fluorescence^34^ and common scoring methods^79^ with our aligned ESM-2 and VAE models with sequences in the reward model test set (Supplementary Table 2). The masked marginal score^7^ is computed with equation (24):

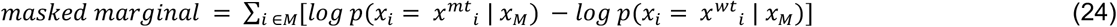

where *M* is the set of mutated positions, *x*^*wt*^ _*i*_ is the parent sequence with no amino acid acid mutations,*x*^*mt*^ _*i*_ is a sequence with amino acid acid mutations at positions *i*, and *x*_*M*_ is a sequence with masks at positions *i* where mutations occur in *x*^*mt*^ relative to *x*^*wt*^. The mutant marginal score^7^ is similarly computed with equation (25):

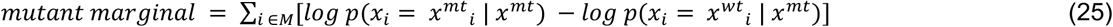

where *x*^*mt*^ is a sequence with amino acids mutations that are not masked. We compute pseudo-perplexity *PP*(*x*)^3,81^ with equation (26):

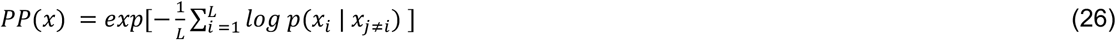

where *L* is the length of a sequence and *p*(*x*_*i*_ | *x*_*j≠i*_) is the probability of amino acid *x* at position *i* given the other amino acids in the sequence *x*_*j≠i*_. For VAE models, We compute the log ratio^45,46^ with equation (27):

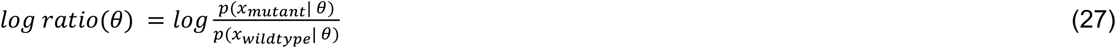

where *x* refers to the mutant or wildtype sequence and *θ* refers to VAE parameters.

### In vivo characterization of sequence variants

To characterize model designs *in vivo*, all designs including wildtype CreiLOV and CreiLOV-T7S, were ordered as clonal pET-28a(+) constructs from Twist Bioscience. Empty pET-28a(+) was also ordered to serve as a baseline. All constructs were transformed into *E. coli* BL21(DE3) competent cells (Thermo Scientific, Catalog No. EC0114) according to a standard heat shock protocol, plated on LB agar plates containing 50 μg/mL of kanamycin sulfate (Sigma-Aldrich, Catalog No. K4000), and grown overnight at 37°C. Single colonies were picked and inoculated in 2 mL starter cultures of LB with 50 μg/mL of kanamycin and grown overnight at 37°C with 225 rpm of orbital shaking. 2 μL of all turbid cultures (including wildtype and empty plasmid) were inoculated in separate wells of a black, optical bottom 96 well plate (Thermo Scientific, Catalog No. 237105) containing 200 μL of LB with 50 μg/mL of kanamycin. The plate was inserted into an Agilent Biotek Synergy H1 plate reader, where the gain and z-position were auto-adjusted (142, 4.75mm respectively), and the excitation and emission bandwidths were set to 9 nm. The plate reader was set to 37°C with orbital shaking set to 237 cpm, and all wells were grown to an OD600nm of approximately 0.5, with A600nm being read every 30 minutes. At the desired confluency, all wells were induced with 0.5 mM of IPTG (Gold Biotechnology, Catalog No. 12481C), and inserted back into the plate reader. The cells were grown for another 210 minutes, with A600 being read every 30 minutes, at which point the plate reader measured the fluorescence at each well using an excitation wavelength of 450 nm and an emission wavelength of 495 nm. We calculated the fluorescence of each design relative to wildtype CreiLOV with equation (28):

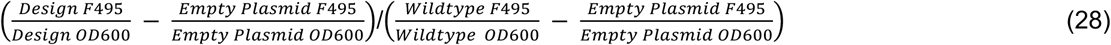

Triplicate biological replicates were performed using the same plate reader parameters as described above to ensure consistency of experimental measurements. The average of these measurements is reported along with the standard error of the mean.

### In vitro characterization of sequence variants

To characterize model designs in vivo, all designs including wildtype CreiLOV and CreiLOV-T7S, were ordered as clonal pET-28a(+) constructs with C-terminal Hisx8 tag from Twist Bioscience. The expression and purification were performed mostly in accordance with method described in Chen2021ACSSynBio. Briefly, CreiLOV constructs were expressed in E. coli BL21(DE3) cells at 37 °C for 3hrs after inducing by adding IPTG to a final concentration of 0.5mM at an optical density at 600nm of 0.4-0.6. After cell lysis, the protein was purified using HisPur NiNTA resin (Thermo Scientific) and buffered exchanged to assay buffer (25mM TrisHCl pH9.0, 1M NaCl).

For quantum yield measurements, 200 μL of protein normalized by A450 was added to a 96-well microplate. The samples were placed in a BioTek Synergy H1 Microplate Reader (Agilent) and illuminated at 450 nm. The emission spectrum between 480 and 610 nm was recorded, subtracted by the average of blank reads, and integrated. The quantum yields were calculated by dividing integrated area by the average of that of WT CreiLOV and then multiplying by the reported WT’s quantum yield value of 0.47.

## Data Availability Statement

The work described uses six existing deep mutational scanning data sets. The first five (GB1, GFP, Ube4B, Bgl3, and Pab1) were retrieved from this prior publication (https://www.pnas.org/doi/10.1073/pnas.2104878118) and the associated GitHub repository (https://github.com/gitter-lab/nn4dms/tree/master/data). The CreiLOV data set was from Data Set S1 and Data Set S2 in the Supporting Information Section of: https://pubs.acs.org/doi/10.1021/acssynbio.2c00662

## Code Availability Statement

All code for running RLXF is available on the public, open-source GitHub repository: https://github.com/RomeroLab/RLXF under the Apache 2.0 license.

