## Supplementary Information for "Functional alignment of protein language models via reinforcement learning"

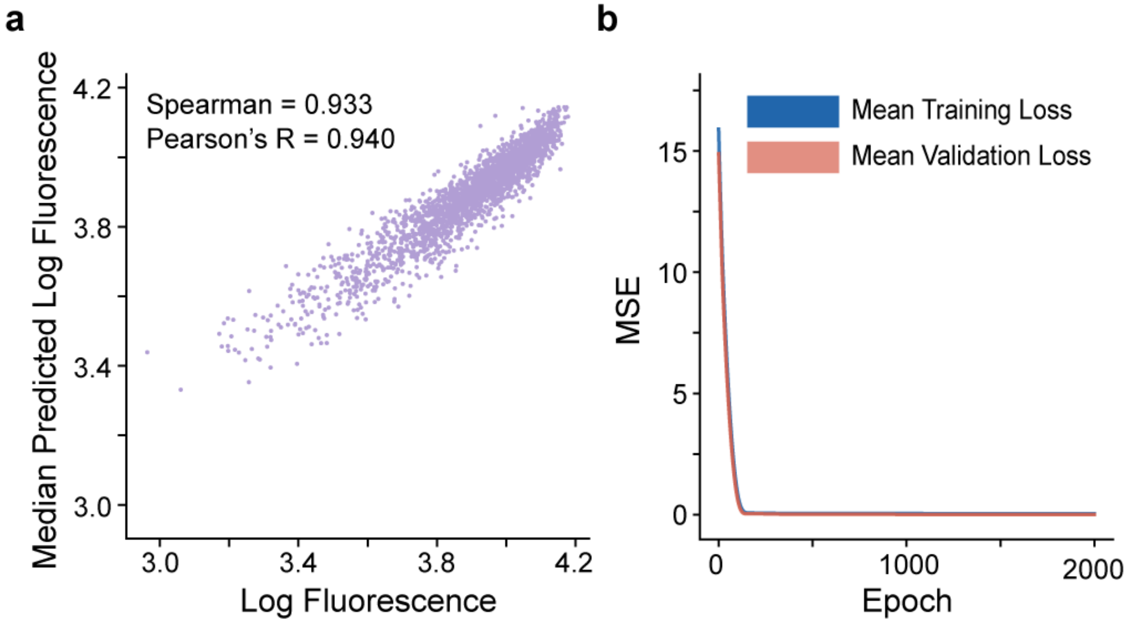


Supplementary Figure 1: Ensemble of reward models for CreiLOV. a, The ensemble of reward models predict the log fluorescence of test set variant sequences with 5 mutations relative to CreiLOV. b, Mean multiple-squared error loss curves across reward model training.

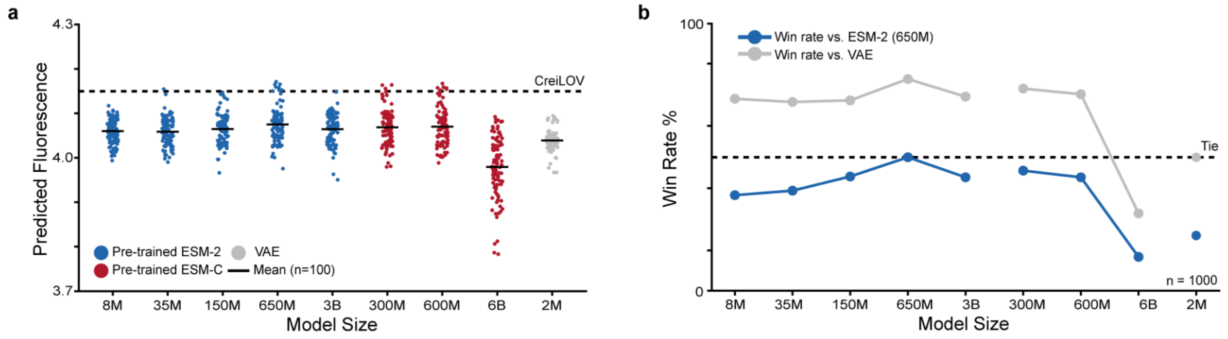


Supplementary Figure 2: Generative capabilities of pre-trained pLMs and our VAE. a, Predicted log fluorescence of sequence variants with 5 mutations relative to CreiLOV sequence from pre-trained ESM-2, ESM-C, and our VAE. b, Win rate of sequence designs from pre-trained ESM-2, ESM-C, and our VAE by random selecting one position to sample and sampling amino acids at that position with highest probability until 5 mutations are introduced. We call this a max sampling strategy.

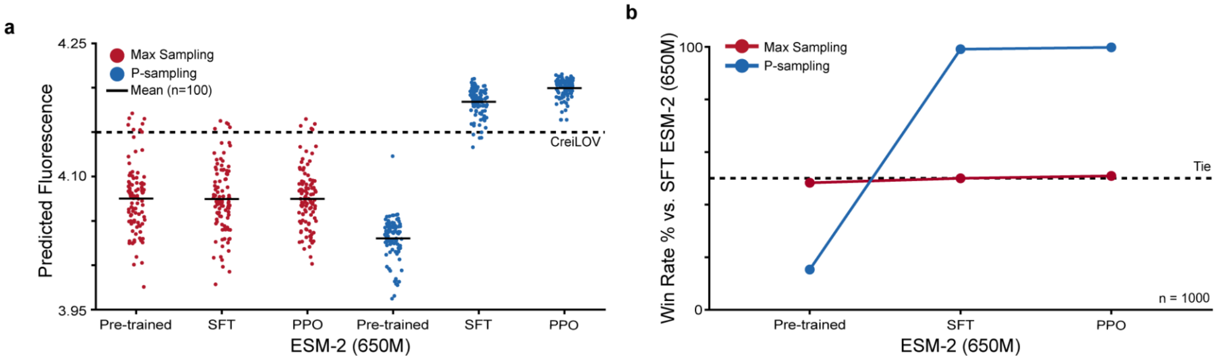


Supplementary Figure 3: Sampling strategies for generation of sequence variants. a, Predicted log fluorescence of sequence designs from ESM-2 (650M) after pre-training, supervised fine-tuning, and proximal policy optimization generated with a max sampling strategy and our top p-sampling strategy. b, Win rate of sequence designs from ESM-2 (650M) after pre-training, supervised fine-tuning, and after proximal policy optimization generated with a max sampling strategy and our p-sampling strategy. Interestingly, we observed that a few mutations significantly enhance predicted fluorescence. Our p-sampling strategy leverages pre-trained ESM-2 (650M) knowledge and knowledge gained during alignment to identify non-natural mutations and sequence positions to mask and sample amino acids that are critical for enhancing function.

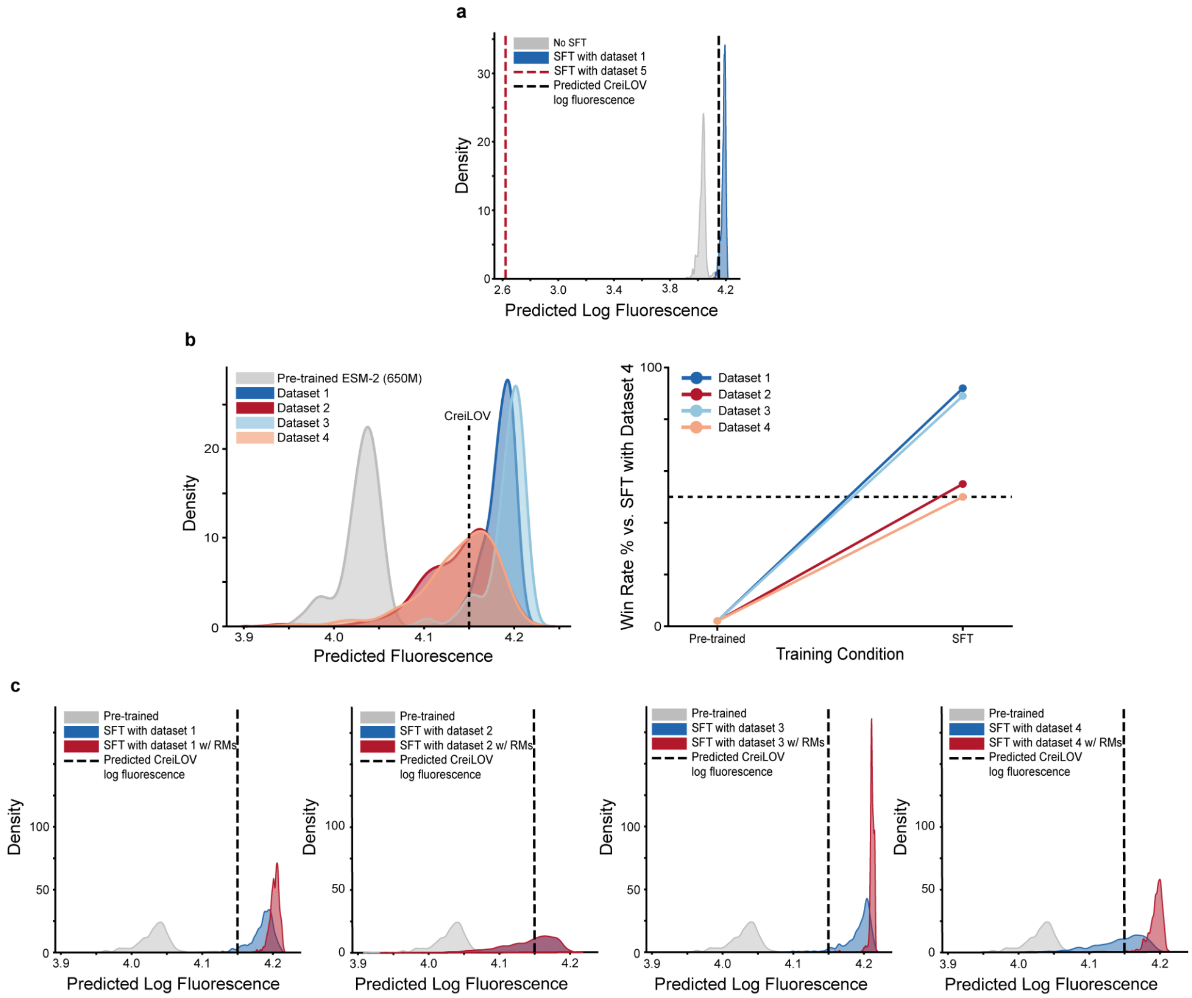


Supplementary Figure 4: Supervised fine-tuning approaches. a, Supervised fine-tuning with synthetic sequence dataset of 5 mutant variants outperforms supervised fine-tuning with a sequence dataset of the brightest variants in the CreiLOV DMS data containing 0-4 mutations relative to CreiLOV (Supplementary Table 1). Interestingly, we observed model collapse/instability during supervised fine-tuning with dataset 5. b, Kernel density estimate plot for predicted log fluorescence and win rate for sequence variants from supervised fine-tuned ESM-2 (650M) models using different synthetic sequence datasets. c, Supervised fine-tuning with synthetic sequence datasets with or without introducing 20 random masks (RMs) to sequence prompts. The distribution of mutations in synthetic sequence datasets are included in Supplementary Figure 5.

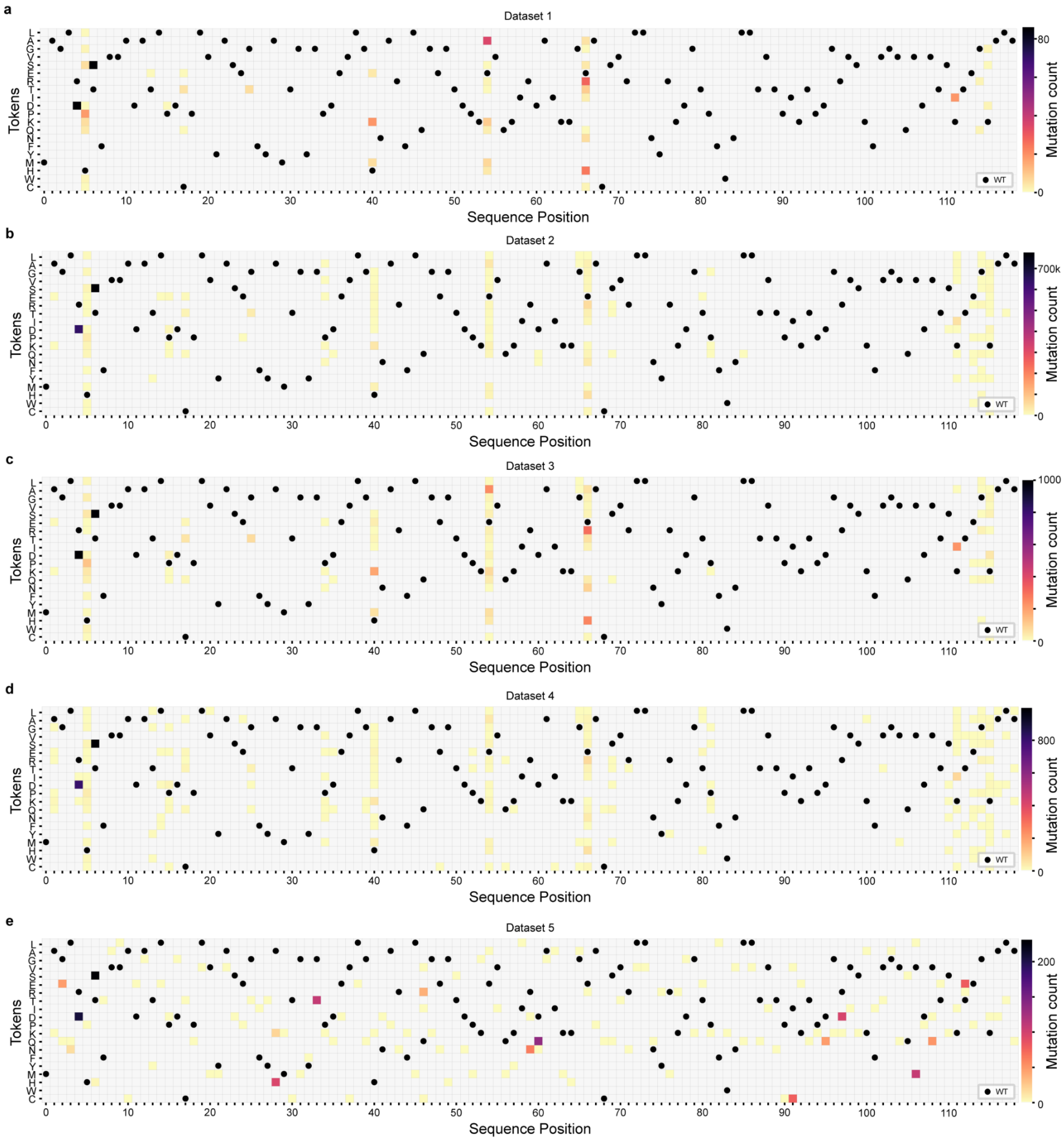


Supplementary Figure 5: Supervised fine-tuning datasets. Mutation count for a, Dataset 1: 86 unique optimized 5 mutant variants of CreiLOV from 100 simulated annealing simulations b, Dataset 2: 780,744 unique 5 mutant variants of CreiLOV saved throughout 100 simulated annealing simulations c, Dataset 3: 1000 unique 5 mutant variants of CreiLOV with the greatest predicted log fluorescence according to ensemble of reward models sampled from dataset 2 d, Dataset 4: 1000 5 mutant variants of CreiLOV randomly sampled from dataset 2 e, Dataset 5: 513 sequence variants of CreiLOV with the best experimental log mean fluorescence and 0-4 mutations in CreiLOV DMS dataset.

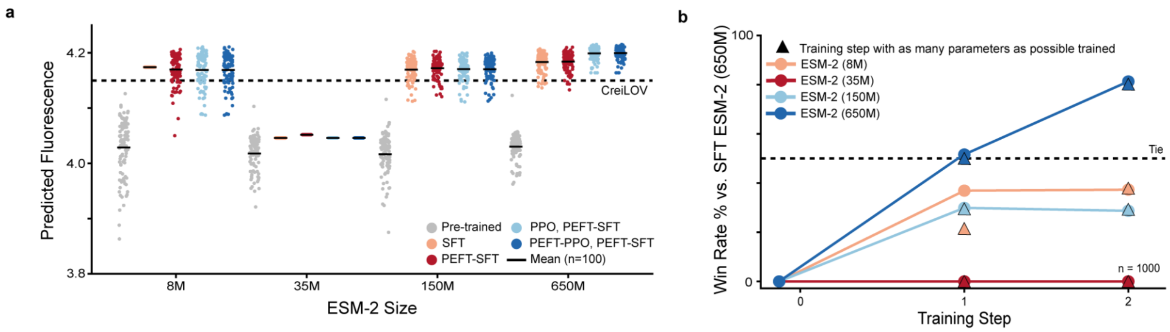


Supplementary Figure 6: ESM parameter-efficient training technique and model scaling effects. a, Predicted log fluorescence of sequence designs from ESM-2 pLMs (8M, 35M, 150M, 650M) after pre-training, supervised fine-tuning, and proximal policy optimization with as many parameters trained as possible on a NVIDIA L40S GPU or our parameter-efficient training technique described in the methods. b, Win rate for sequence designs from ESM-2 pLMs (8M, 35M, 150M, 650M) after pre-training, supervised fine-tuning, and proximal policy optimization with as many parameters trained as possible on a NVIDIA L40S GPU or our parameter-efficient training technique described in the methods.

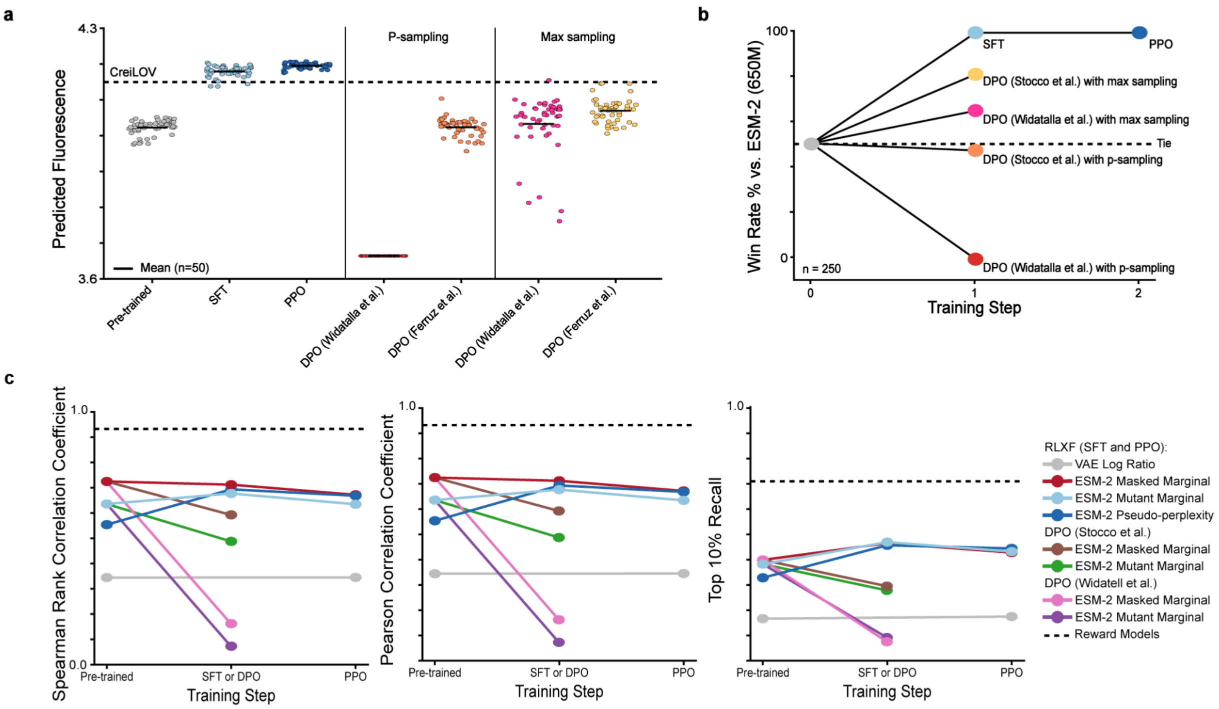


Supplementary Figure 7: Benchmarking RLXF against related methods. a, Predicted log fluorescence of designs from ESM-2 (650M) aligned via RLXF or DPO. We used our p-sampling strategy for models aligned with RLXF. Models fine-tuned with DPO generated more fluorescent sequence variants when generating sequence variants via a max sampling strategy instead of our p-sampling strategy. b, Win rate for designs from models aligned via RLXF or DPO. c, Common metrics for ESM-2 (650M) and our VAE during alignment with RLXF or DPO. Interestingly, zero-shot predictions often decreased after RLXF or DPO except for top 10% recall. Top 10% recall is likely a better metric for estimating the generative capabilities of a pLM.

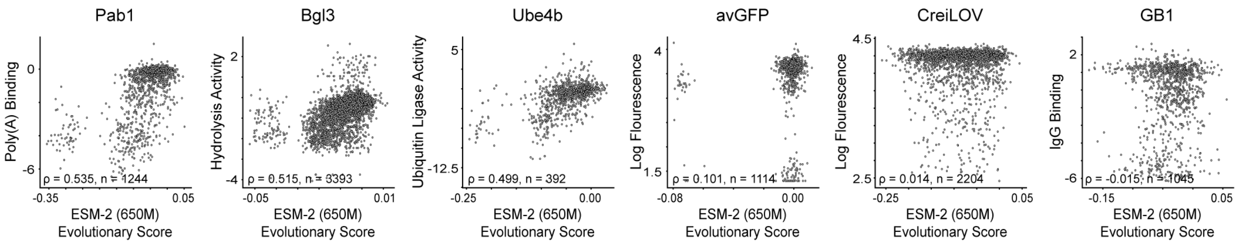


Supplementary Figure 8: Deep mutational scanning (DMS) measurements vs. ESM-2 (650M) evolutionary scores. For each protein, we compare experimental DMS readouts to an ESM-2 (650M) evolutionary score. Scores were computed using a fast WT-logits pseudo-likelihood approach. We pre-computed ESM-2 logits for the wild-type sequence with each position masked once, then for every variant we read off the log-probability of its residue at each WT-aligned position under those masked-WT distributions. We summarize this as a per-sequence mean negative log-likelihood and report a WT-normalized score (greater than 0 indicates a sequence variants is ranked higher than WT). Each plot reports the Spearman's correlation and the number of scored sequence variants. Sequence variants here each contain 1 mutation (Supplementary Table 2).

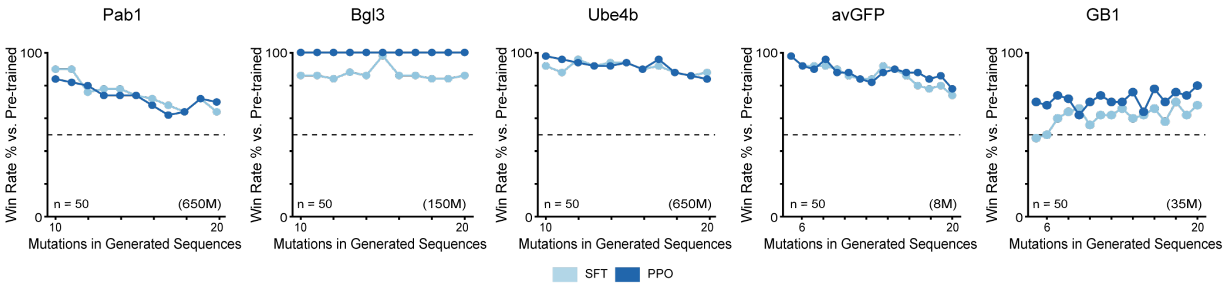


Supplementary Figure 9: Win rate for sequence designs with varying number of mutations from SFT ESM-2 and PPO ESM-2 relative to pre-trained ESM-2. We used the ESM-2 model size with the top PPO win rate from panel Figure 2a (model size indicated in parentheses).

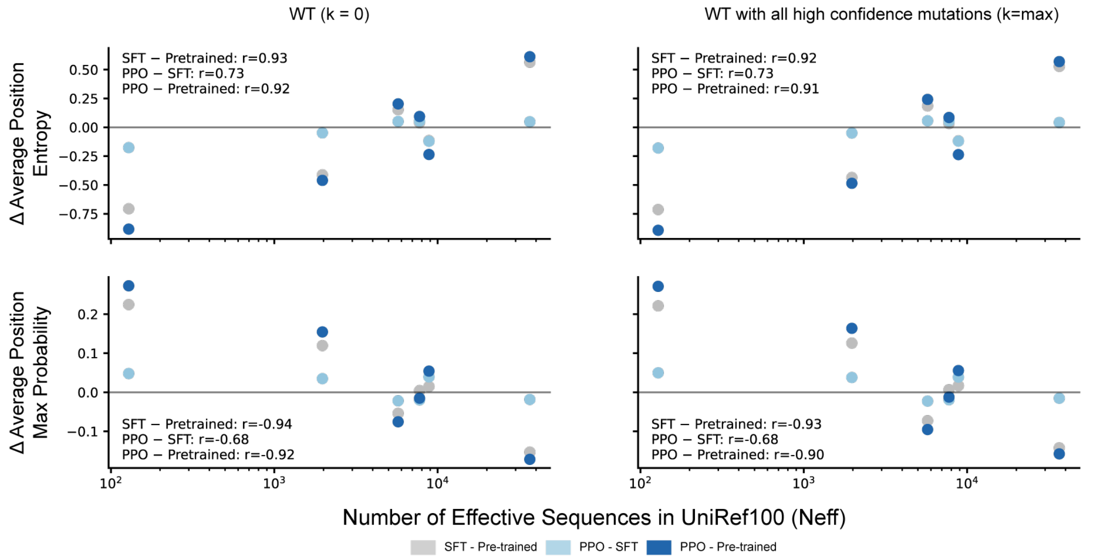


Supplementary Figure 10: Summary statistics of single-mutant logits across alignment stages for wildtype and high-confidence mutant sequences. For each benchmark protein, we report the change in the average positional entropy and max probability from ESM-2 single-mutant logits computed on the WT sequence and along a high-confidence mutation trajectory, illustrating how SFT and PPO-based RLXF systematically shift and/or rescale local residue preferences relative to the pre-trained prior.

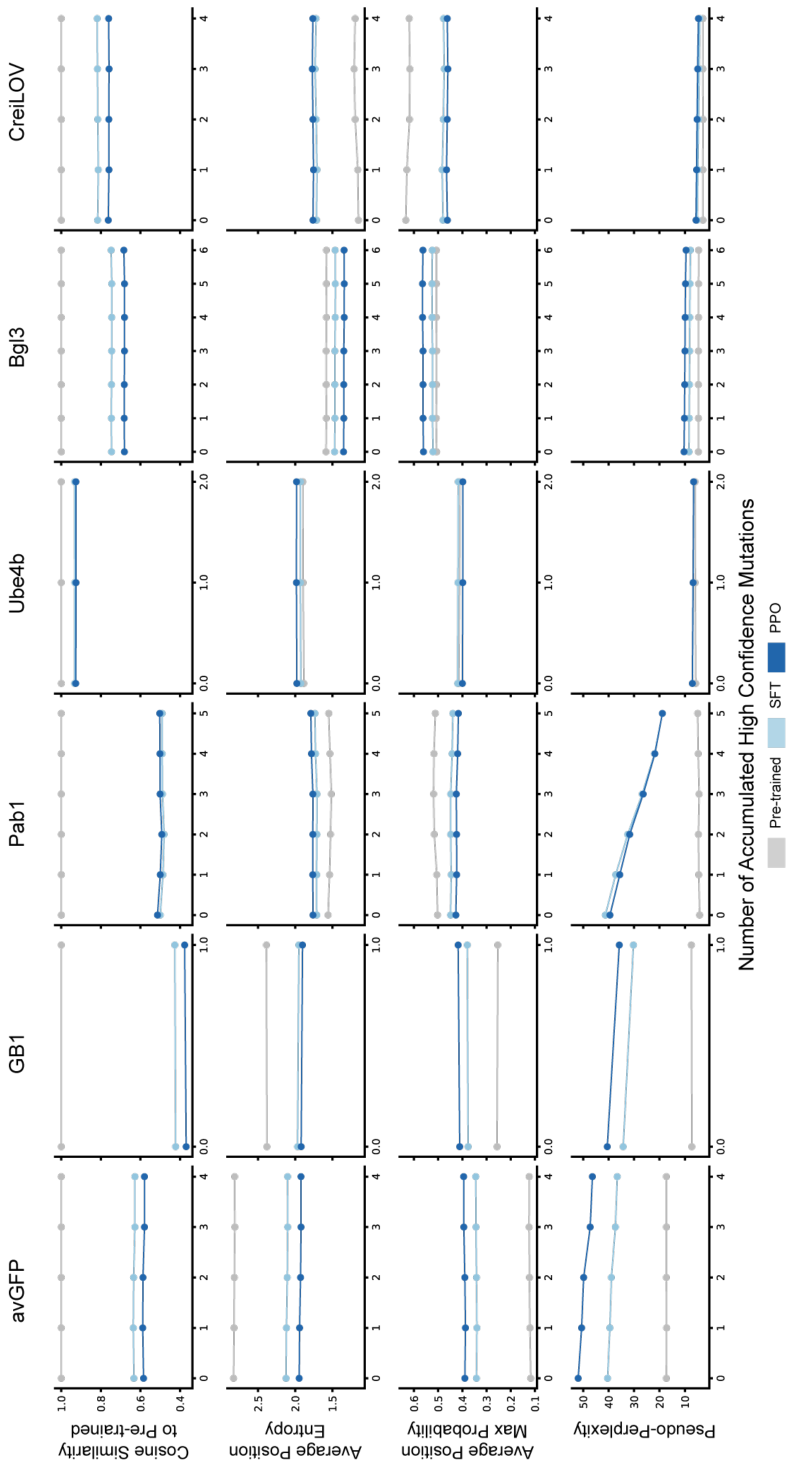


Supplementary Figure 11: Distribution of single-mutant logits for the wildtype sequence and high-confidence mutant sequences. For each benchmark protein, we compare the distribution of ESM-2 logits for the wildtype sequence (WT) against logits obtained along a high-confidence mutation trajectory, highlighting how supervised fine-tuning (SFT) and PPO-based RLXF shift local residue preferences relative to the pre-trained prior.

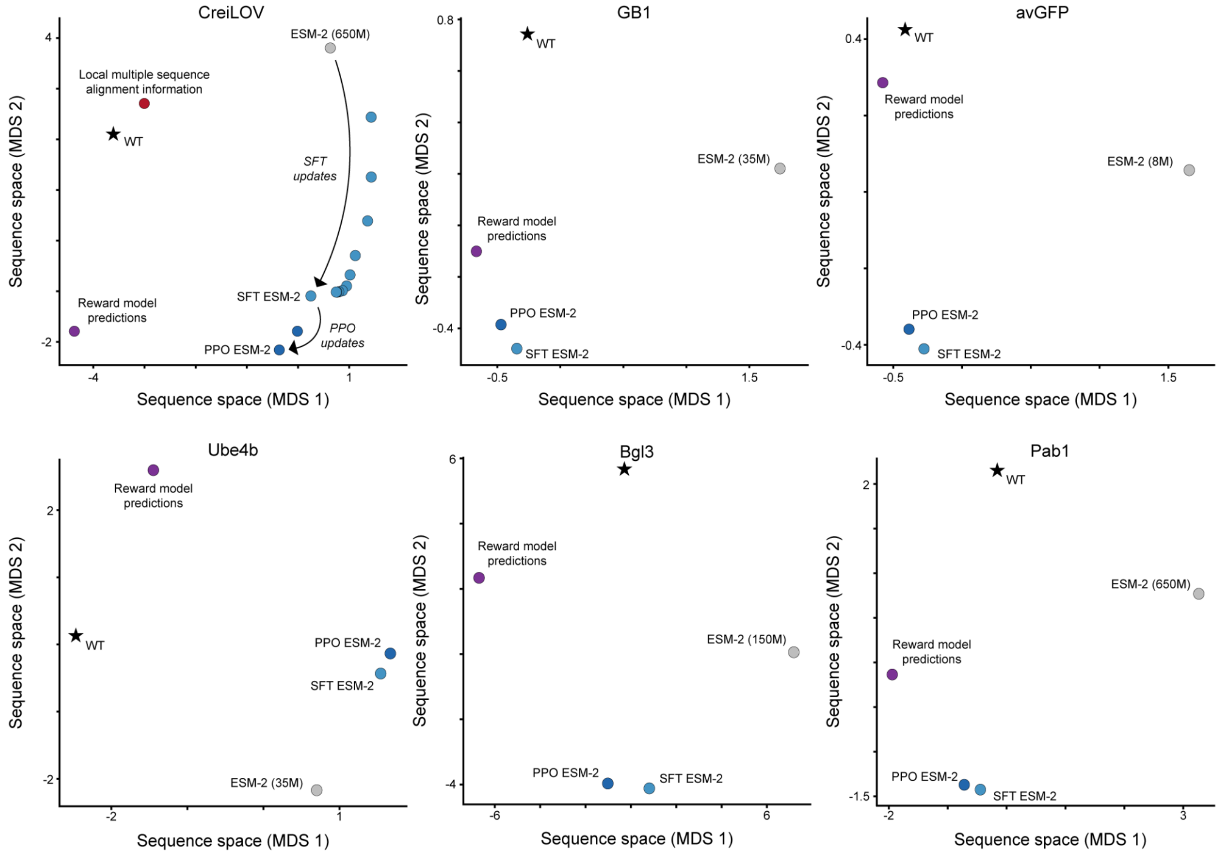


Supplementary Figure 12: Multi-dimensional scaling plots of Euclidean distance for single mutation probability matrices derived from different models. Pre-trained ESM-2 initially generated sequences that occupied a distinct sequence space but learned mutational preferences more consistent with reward models trained on experimental deep mutational scanning (DMS) data after supervised fine-tuning (SFT) and proximal policy optimization (PPO). The natural wildtype sequence (WT) was represented with a one-hot encoded matrix. ESM-2 probabilities were generated by masking the natural wildtype sequence (WT) at one position at a time and applying the softmax to the output logits produced. Reward models predicted the function of all single mutants for the natural wildtype sequence (WT) and assembled into a matrix indexed by position and amino acid mutation to represent reward model predictions. We performed linear regression to account for differences in scale between matrices. Euclidean distances between the regression-predicted matrices using pre-trained ESM-2 as the target matrix were computed, and multi-dimensional scaling (MDS) was performed to visualize the relationships. For CreiLOV, VAE probabilities were also obtained by inputting the natural wildtype sequence for each protein into the VAE and taking the softmax over the output logits at each position. We considered this local multiple sequence alignment information.

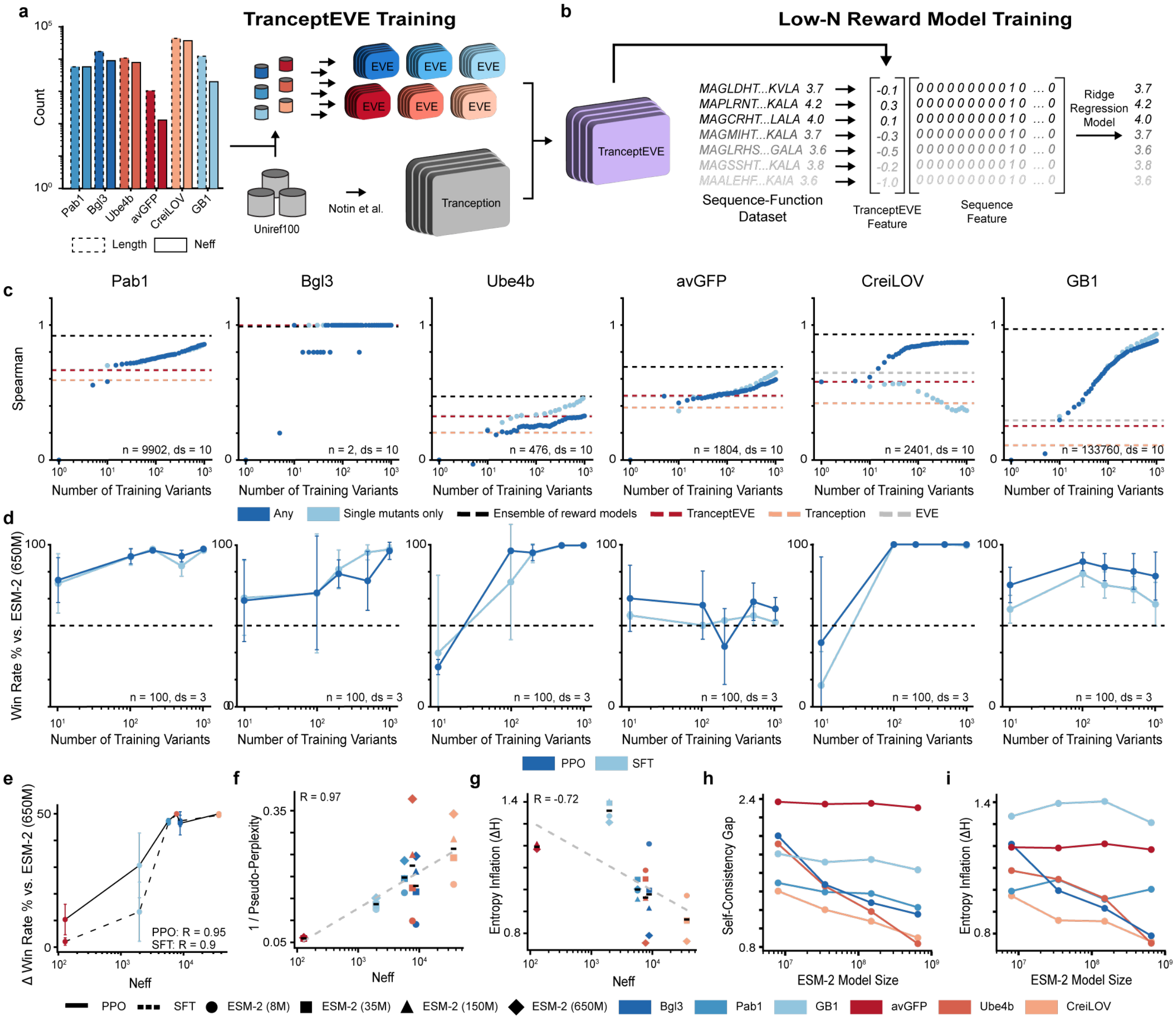

Supplementary Figure 13: Low-N functional alignment reveals importance of well-calibrated protein language model priors. a, TranceptEVE training. b, Low-N reward modeling training. c, Spearman correlation for test set sequences from Supplementary Table 2 for ridge regression reward models averaged across 10 training data splits (ds) for training set sizes up to 1000 sequence variants. d, Win rate of sequence designs from pre-trained, SFT, and PPO ESM-2 relative to base model ESM-2 (650M). e, Change in win rate for sequence designs after SFT and PPO relative to base model ESM-2 (650M) and number of effective sequences (Neff) from UniRef100 for each protein. f, Inverse pseudo-perplexity from pre-trained ESM-2 (650M) and number of effective sequences (Neff) from UniRef100 for each protein and ESM-2 model size. We indicate inverse pseudo-perplexity averaged across model sizes with grey dashed line. g, Entropy inflation for log probabilities from pre-trained ESM-2 (650M) for a position before and after masking a position averaged across each position. We indicate entropy inflation averaged across model sizes with a grey dashed line. h, Self-consistency gap for log probabilities for each model size of pre-trained ESM-2 for a position before and after masking a position averaged across each position. i, Entropy inflation for log probabilities for each model size of pre-trained ESM-2 for a position before and after masking a position averaged across each position.

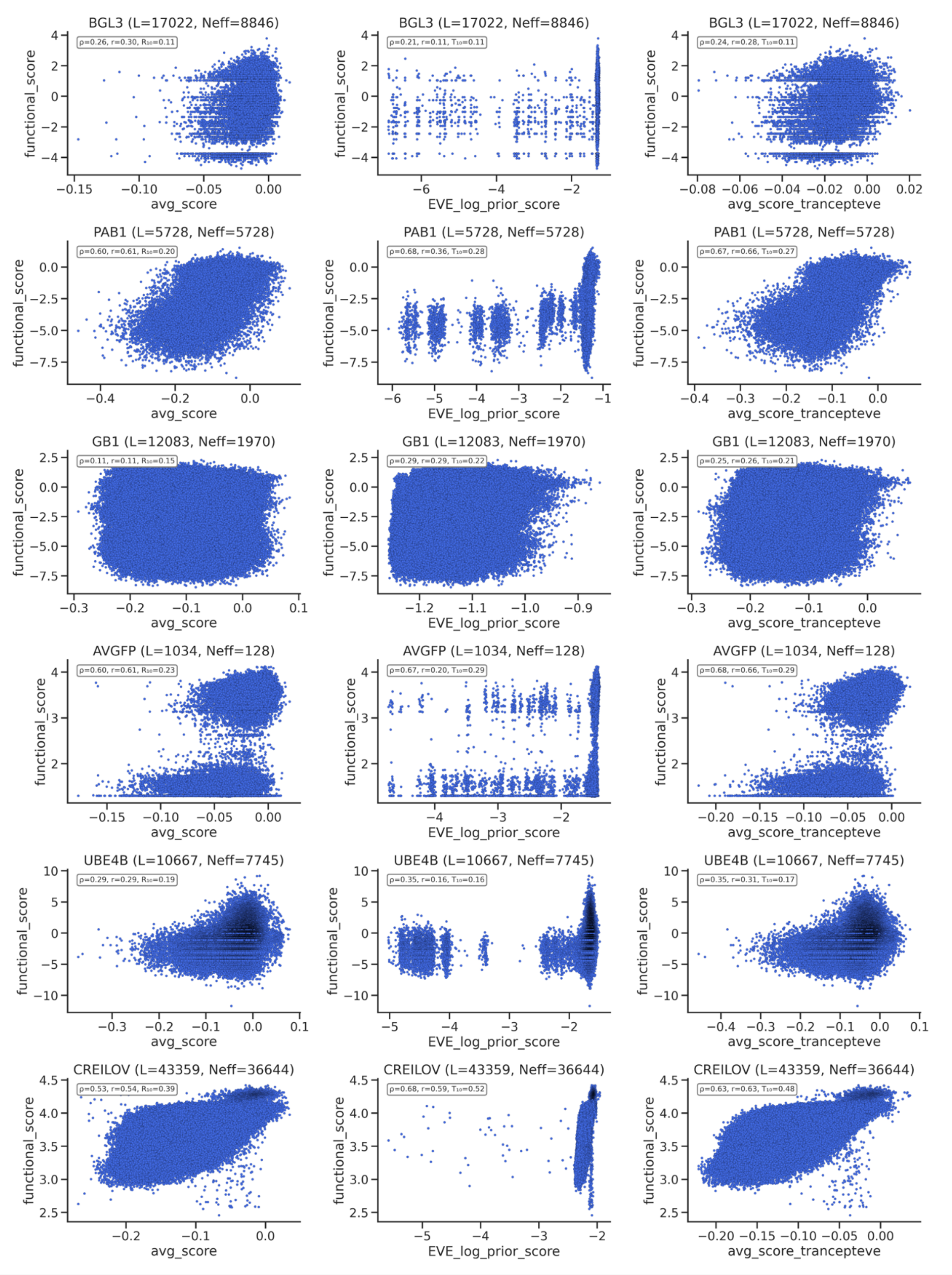


Supplementary Figure 14: Zero-shot Spearman, Pearson, and top 10% recall correlations for Tranception, EVE, and TranceptEVE models for data splits from Supplementary Table 2.

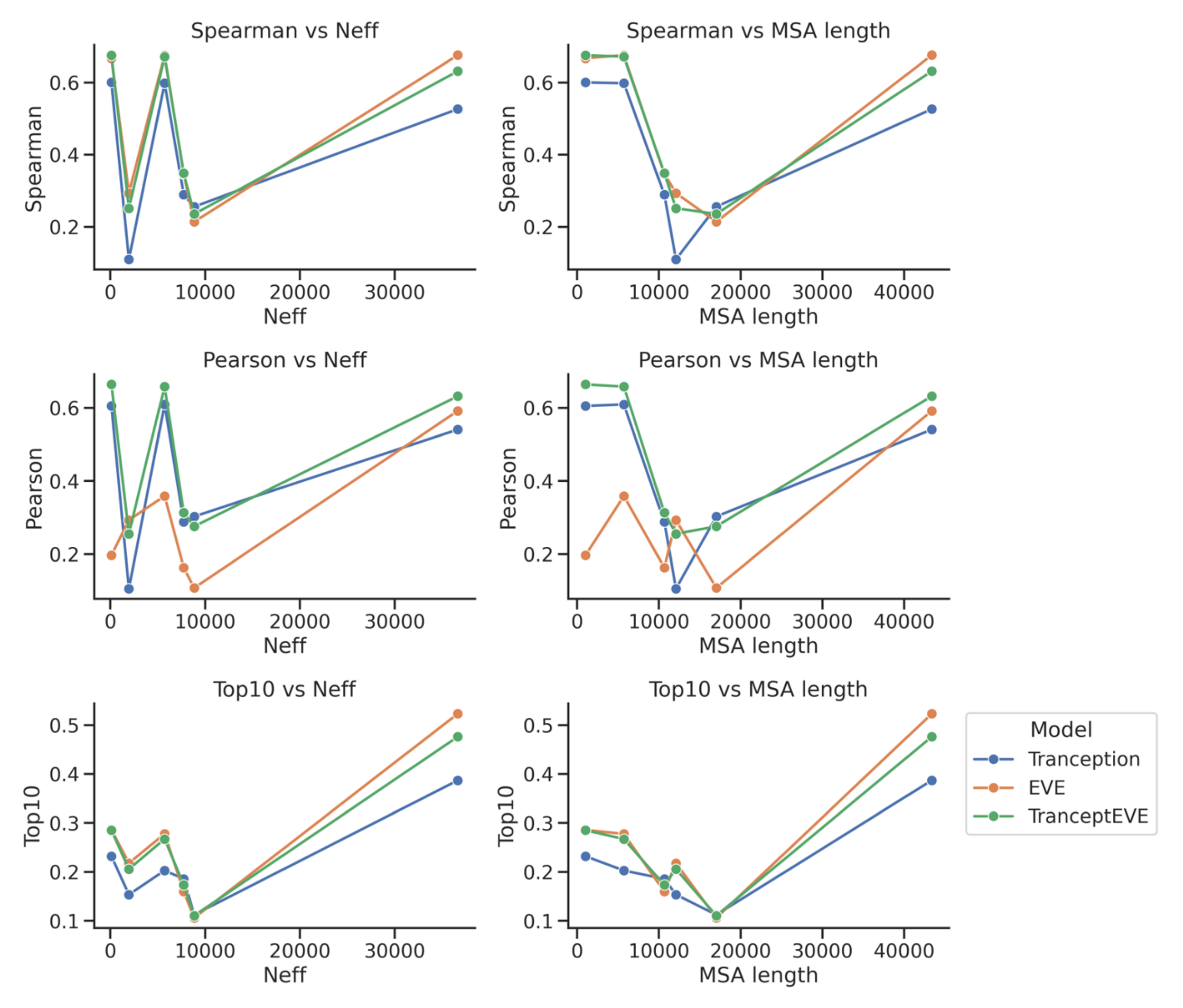


Supplementary Figure 15: Summary of zero-shot Spearman, Pearson, and top 10% recall correlations for Tranception, EVE, and TranceptEVE models with MSA effective size (Neff) and MSA length for all data splits from Supplementary Table 2. There was no obvious relationship.

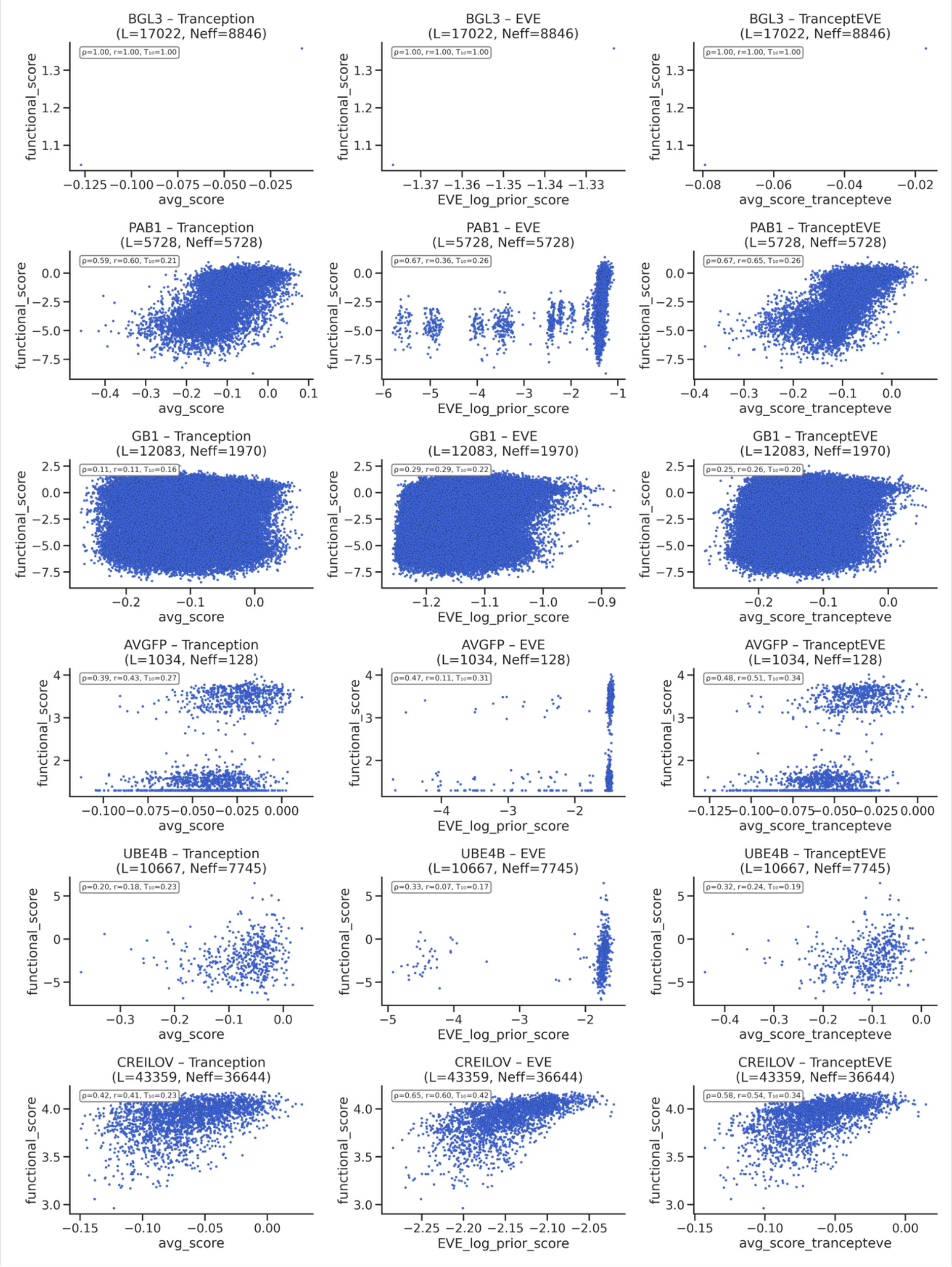


Supplementary Figure 16: Zero-shot Spearman, Pearson, and top 10% recall correlations for Tranception, EVE, and TranceptEVE models for test data split only from Supplementary Table 2.

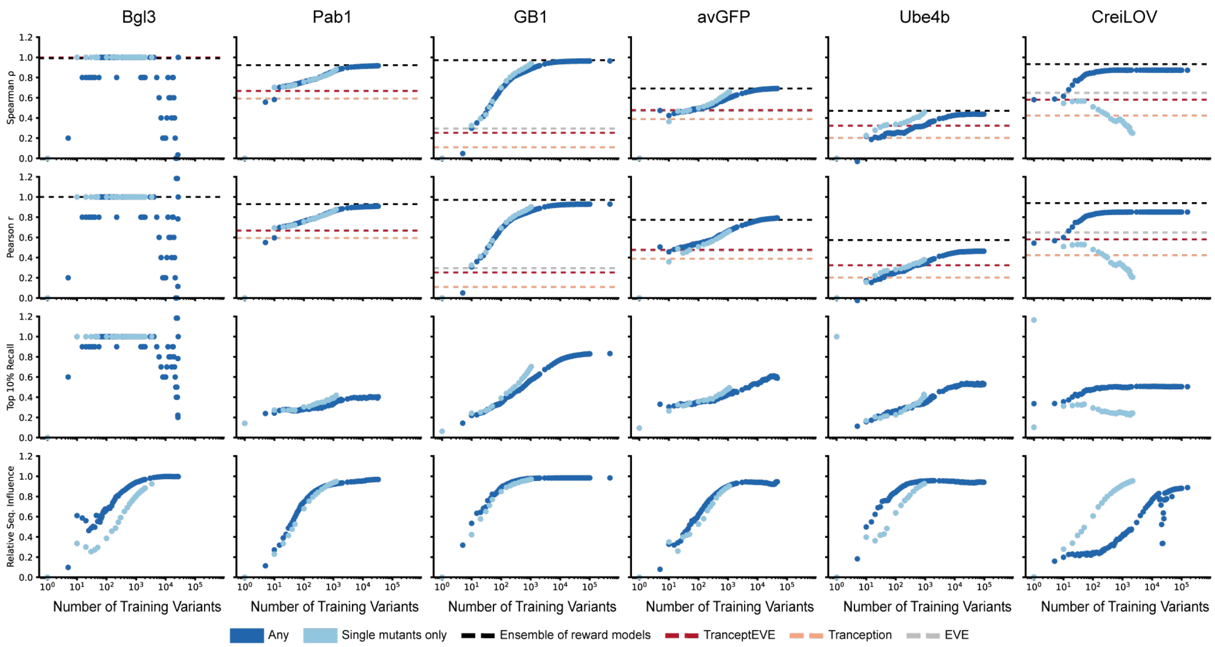


Supplementary Figure 17: Spearman rank correlation, Pearson correlation, Top 10% recall, and Relative Sequence Influence for test set sequences from Supplementary Table 2 for ridge regression reward models averaged across 10 training data splits (ds) for training set sizes up to the max number of training sequence variants. Relative sequence influence was calculated using the sum of weights for one-hot encoded sequence features relative to the sum of weights for one-hot encoded sequence features and the TranceptEVE evolutionary density feature. Low-N reward models trained on fewer than 1,000 variants consistently improved test-set Spearman correlations beyond the strong TranceptEVE zero-shot baseline and in some cases approached the performance of reward-model ensembles trained on substantially larger datasets. Predictive performance was similar whether reward models were trained on combinatorial or single-mutant variants for all proteins besides CreiLOV. The predictive performance for CreiLOV variants degraded with increasing training set size when training exclusively on single mutants, suggesting the strong presence of epistasis in the sequence to function mapping.

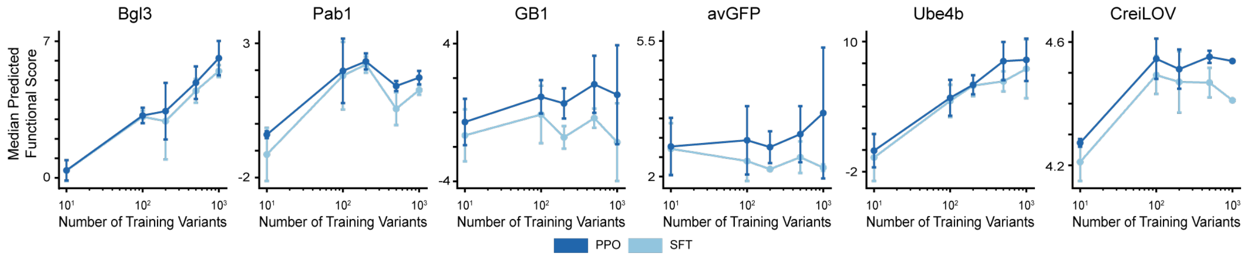


Supplementary Figure 18: Low-N augmented ridge reward models guide functional alignment across protein families. For each benchmark protein and training-set size, we quantify how low-data augmented ridge regression reward models steer RLXF by tracking the predicted fitness of sequences sampled from the pre-trained, SFT, and PPO policies. Curves report the average predicted mean/median/max reward according to low-N ridge regression reward models (error bars indicate variability across 3 replicates), demonstrating that ridge-based reward signals remain sufficient to drive consistent improvements in generation quality under data scarcity.

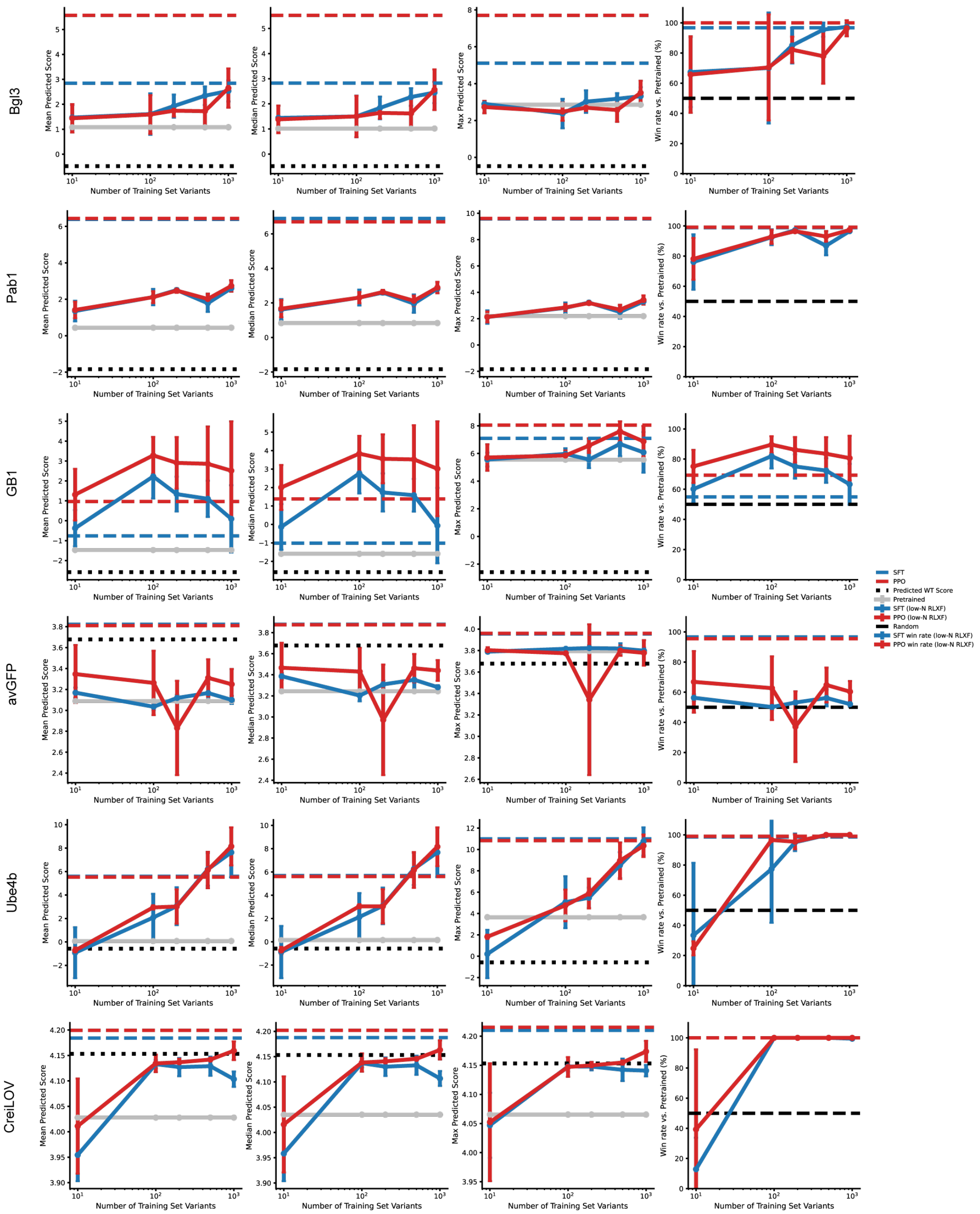


Supplementary Figure 19: Low-N functional alignment with RLXF improves sequence generation across diverse protein classes. For each benchmark protein and training-set size, we report the average mean, median, and maximum predicted reward and win rate for sequences generated at each alignment stage (pre-trained, SFT, PPO) from ESM-2 (650M). Models were aligned using low-N augmented ridge reward models, while these reported scores and win rates were computed using the ensemble of reward models trained on the complete training set, with error bars indicating variability across 3 replicates.

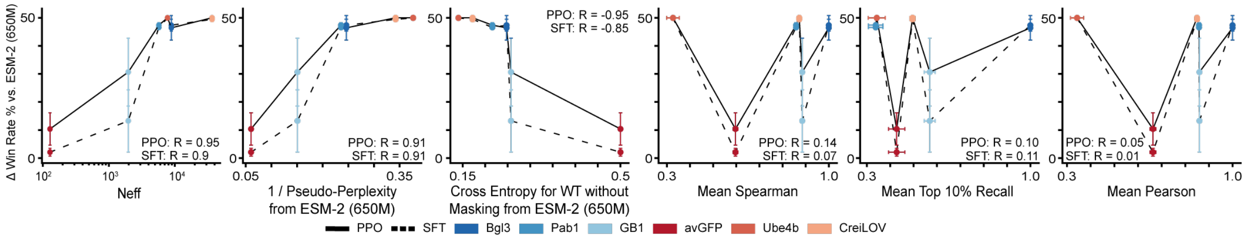


Supplementary Figure 20: Low-N alignment gains rely on strong evolutionary priors and less on predictive performance of low-N ridge regression reward models. For each benchmark protein and training-set size, we relate improvements in generation quality after SFT and PPO to the predictive performance of the corresponding low-N augmented ridge regression reward models. Points denote independent training replicates, showing that reward-model accuracy alone does not reliably predict alignment outcomes, consistent with low-N RLXF depending strongly on the quality of the pre-trained evolutionary prior.

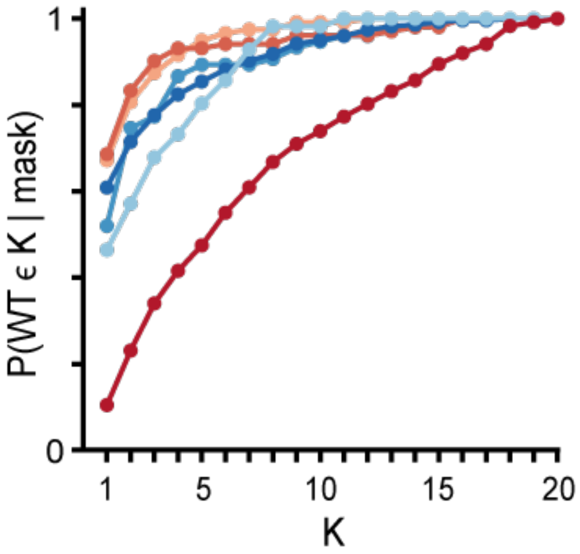


Supplementary Figure 21: Wildtype residue retention under masked inference varies across protein families for ESM-2 (650M). For each benchmark protein, we quantify prior sharpness for the wildtype sequence by measuring the probability assigned to the wildtype amino acid when each position is masked, averaged across sequence positions. Trends across proteins highlight systematic differences in residue-level calibration of pre-trained priors, with deeper-evolution families exhibiting higher wildtype residue probability under masking. Notably, ESM-2 does not assign substantial probabilities to avGFP residues during masked inference.

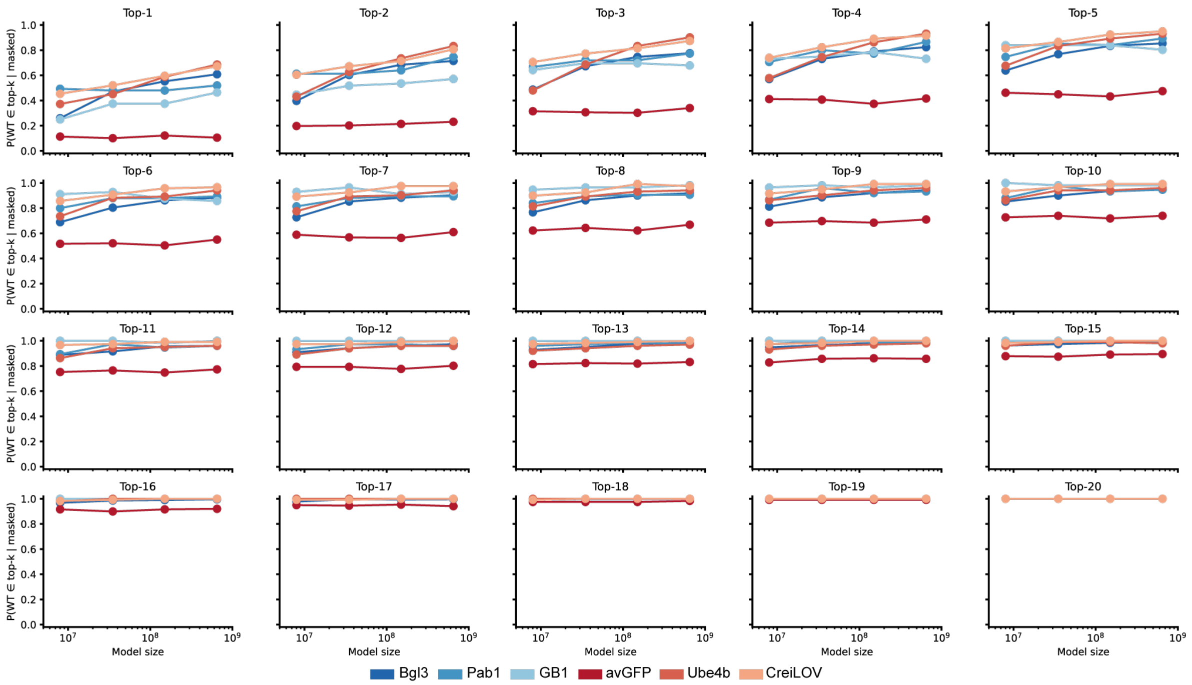


Supplementary Figure 22: Wildtype residue retention under masked inference varies across protein families and model sizes. For each benchmark protein and ESM-2 model size, we quantify prior sharpness for the wildtype sequence by measuring the probability assigned to the wildtype amino acid when each position is masked, averaged across sequence positions. Trends across proteins and sizes highlight systematic differences in residue-level calibration of pre-trained priors, with deeper-evolution families exhibiting higher wildtype residue probability under masking. Notably, ESM-2 does not assign substantial probabilities to avGFP residues during masked inference.

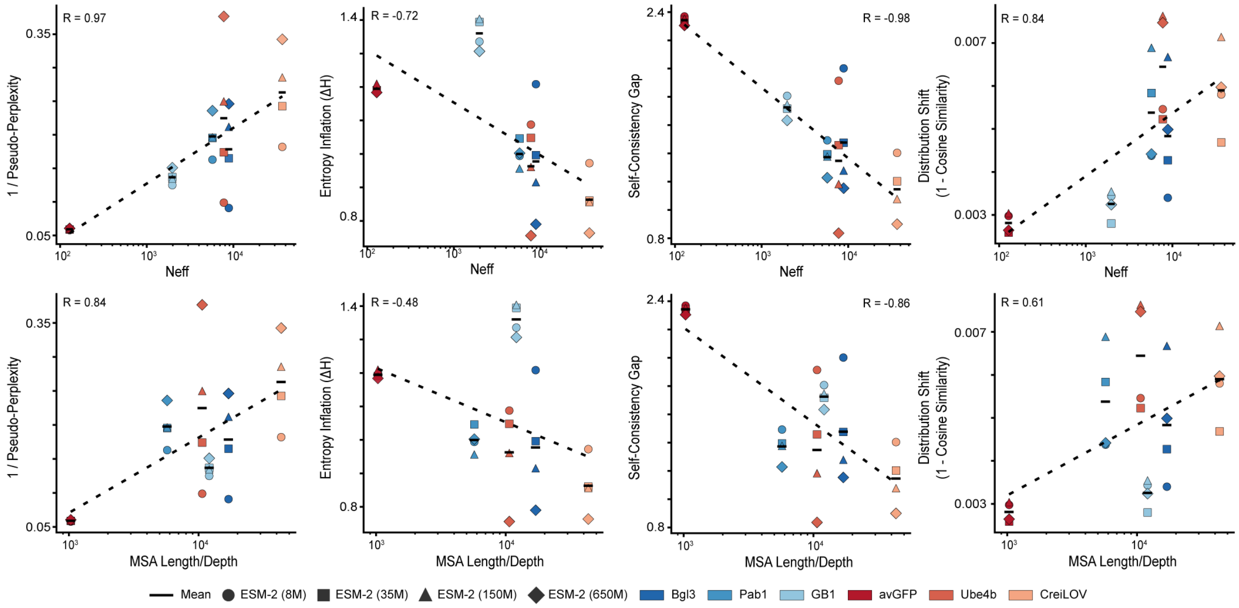


Supplementary Figure 23: Pre-trained ESM-2 priors correlate with evolutionary depth across protein families. For each benchmark protein and ESM-2 model size, we relate evolutionary depth measured by Neff (top row) or MSA depth (bottom row) to prior metrics computed on the wildtype sequence including inverse pseudo-perplexity and metrics calculated comparing logits before and after masking including entropy inflation (ΔH), self-consistency gap (SCG), and distribution shift (1 − cosine similarity). Pearson correlations across proteins highlight that deeper and more diverse MSAs are associated with stronger and more stable pre-trained priors.

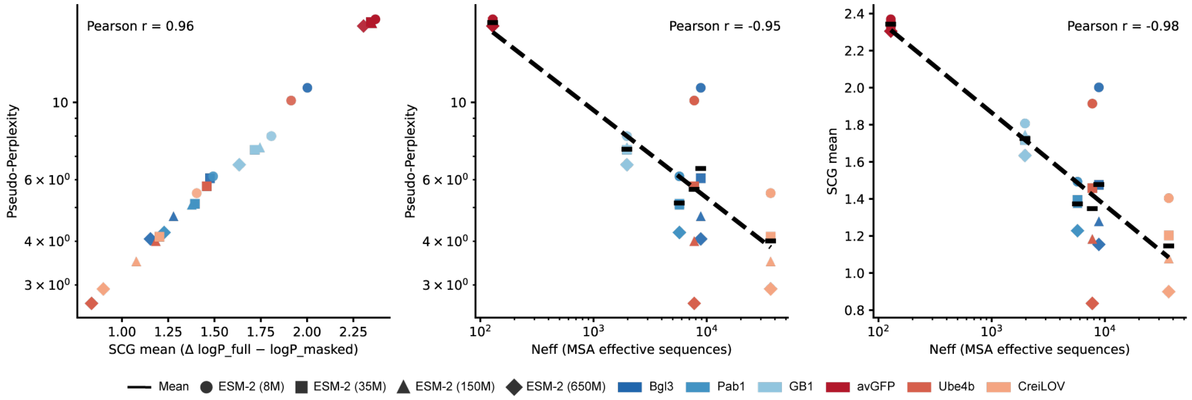


Supplementary Figure 24: Inverse pseudo-perplexity tracks evolutionary depth and prior calibration across protein families. For each benchmark protein and ESM-2 model size, we relate inverse pseudo-perplexity of the wildtype sequence to measures of evolutionary depth (Neff) and additional prior-quality metrics derived from masked inference. Points denote individual proteins and model sizes, highlighting that families with deeper and more diverse MSAs exhibit lower pseudo-perplexities and more well-calibrated pre-trained priors.

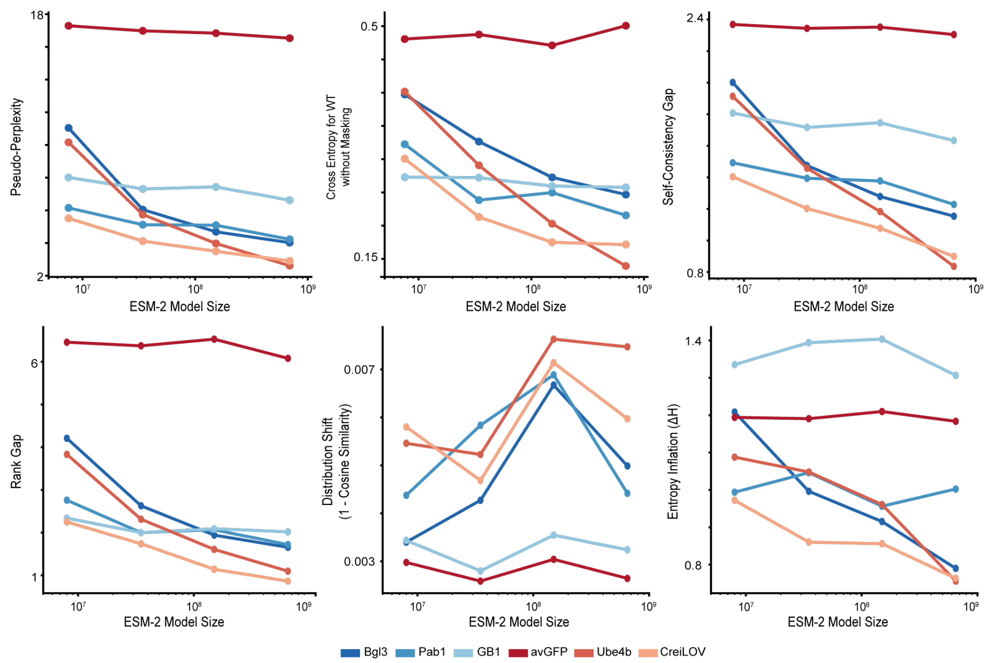


Supplementary Figure 25: Masking sensitivity reveals protein-dependent calibration differences in pre-trained ESM-2 priors across model sizes. For each benchmark protein and ESM-2 model size, we quantify how model predictions change under masked inference using multiple prior-calibration metrics computed on the wildtype sequence including pseudo-perplexity, unmasked wildtype cross entropy, self-consistency gap (SCG), wildtype rank gap, distribution shift ($1 - $ cosine similarity), and entropy inflation ($\Delta H$). Metrics are averaged across sequence positions and summarize changes in log-probability statistics before versus after masking. Protein-specific trends across model sizes highlight systematic variation in prior stability, consistent with weaker calibration in families with shallower evolutionary support.

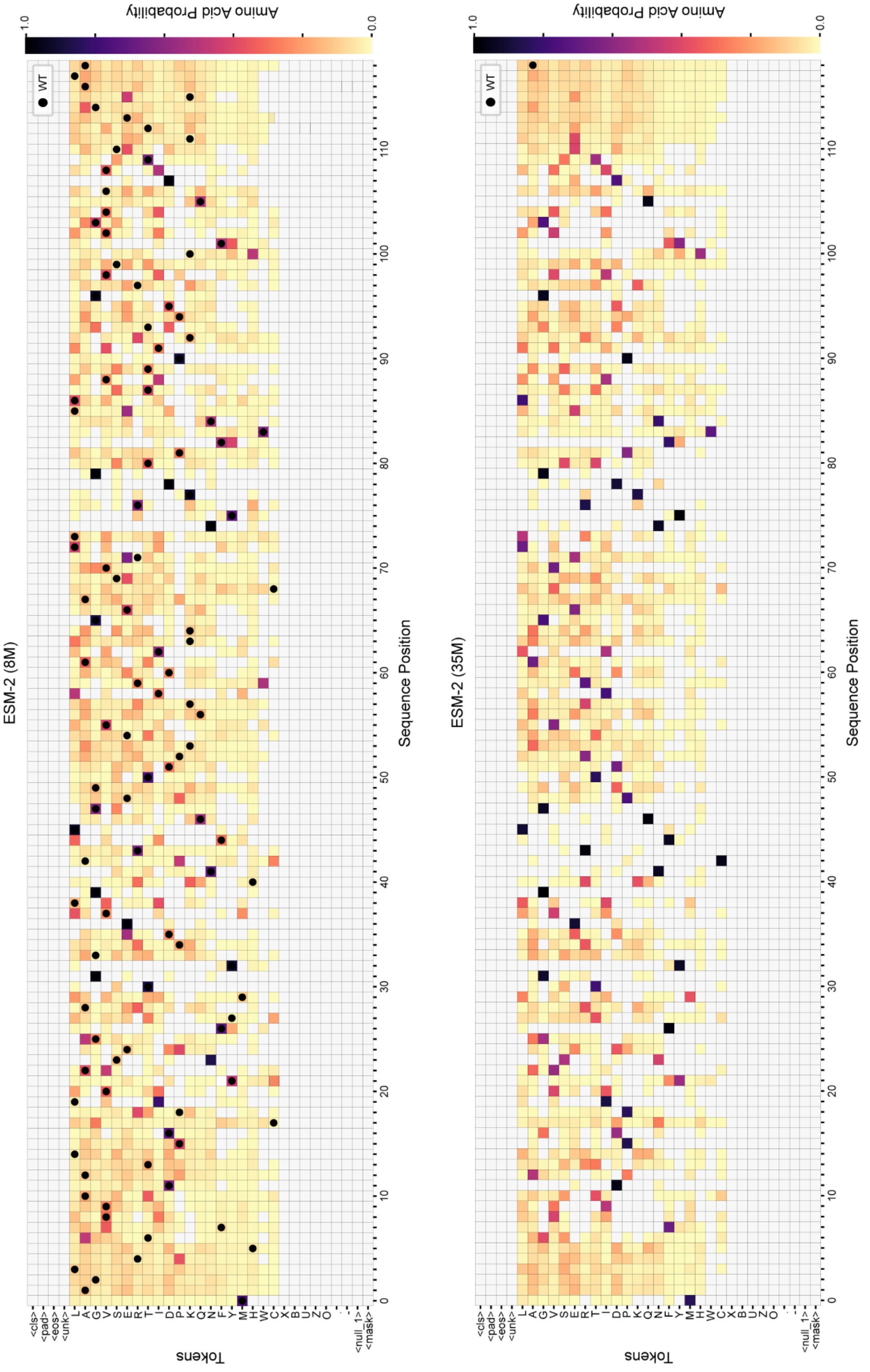


Supplementary Figure 26: Single mutant probabilities for CreiLOV sequence from ESM-2 (8M) and ESM-2 (35M) models. Single mutant probabilities were obtained by masking CreiLOV one position at a time and taking the softmax of logits generated by ESM-2 models.

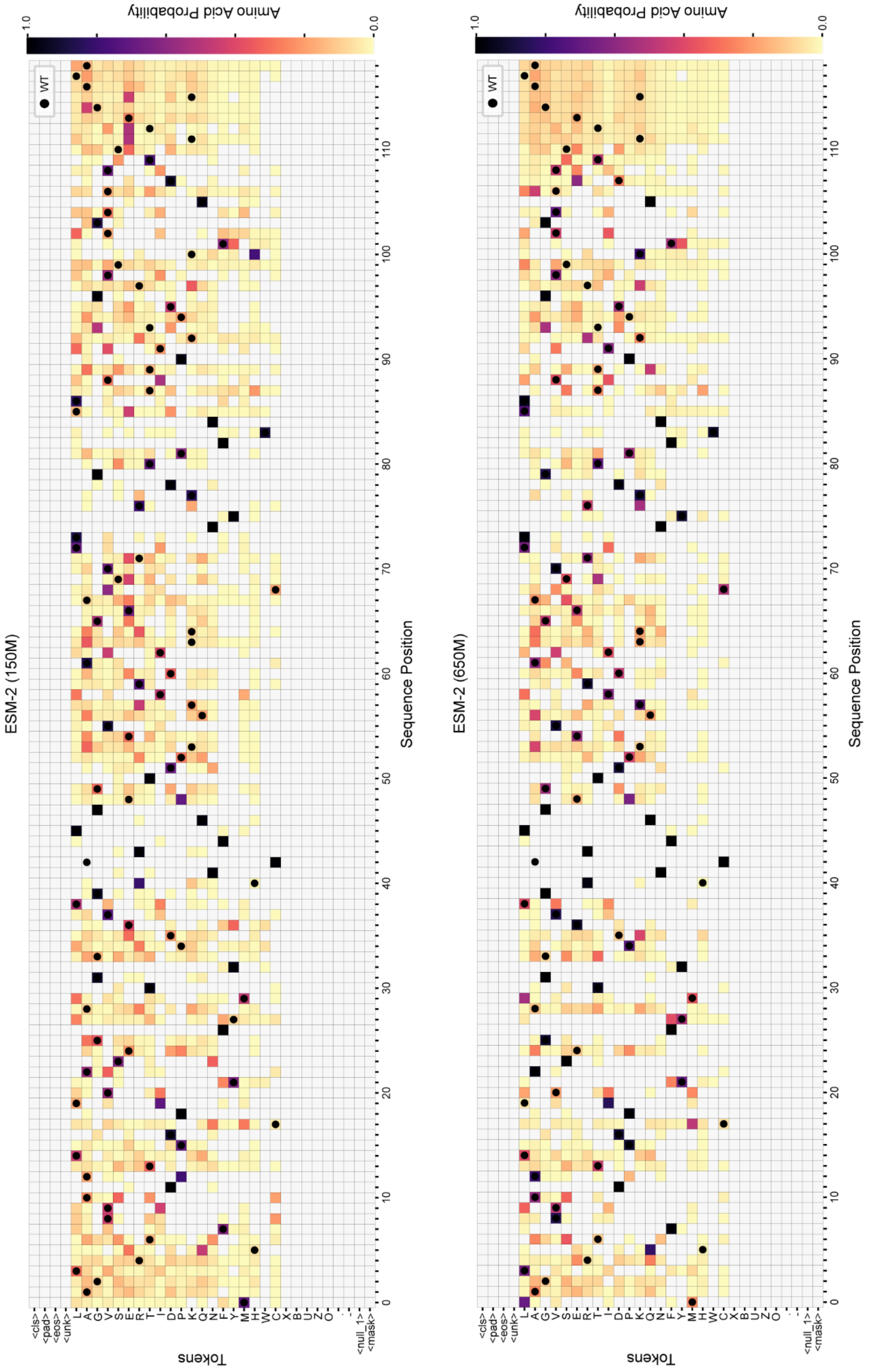


Supplementary Figure 27: Single mutant probabilities for CreiLOV sequence from ESM-2 (150M) and ESM-2 (650M) models. Single mutant probabilities were obtained by masking CreiLOV one position at a time and taking the softmax of logits generated by ESM-2 models.

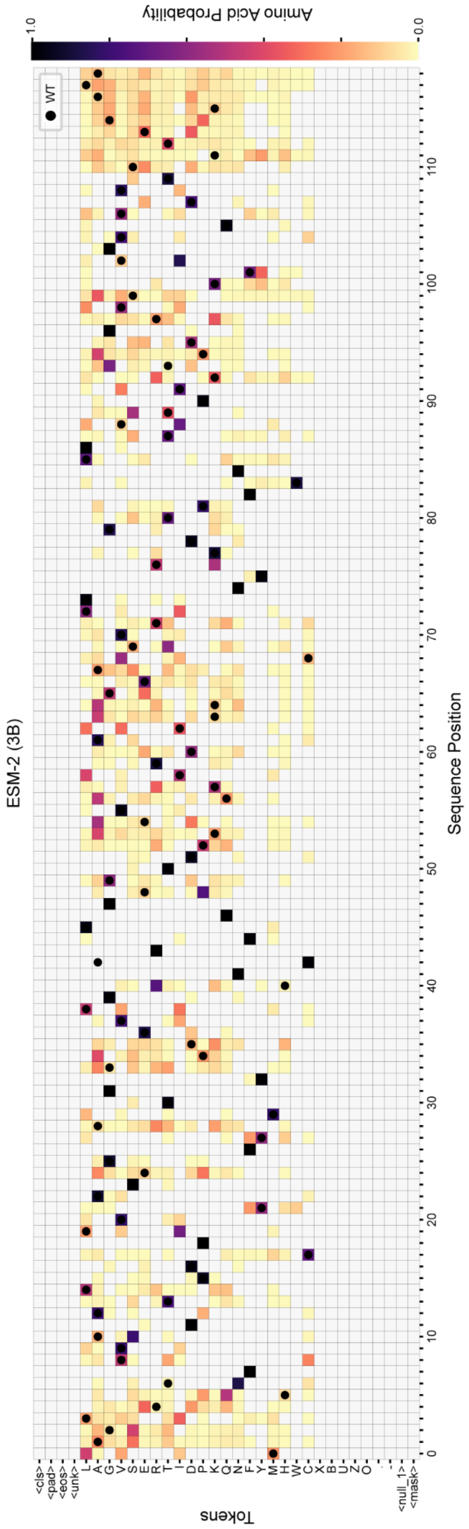


Supplementary Figure 28: Single mutant probabilities for CreiLOV sequence from ESM-2 (3B). Single mutant probabilities were obtained by masking CreiLOV one position at a time and taking the softmax of logits generated by ESM-2 models.

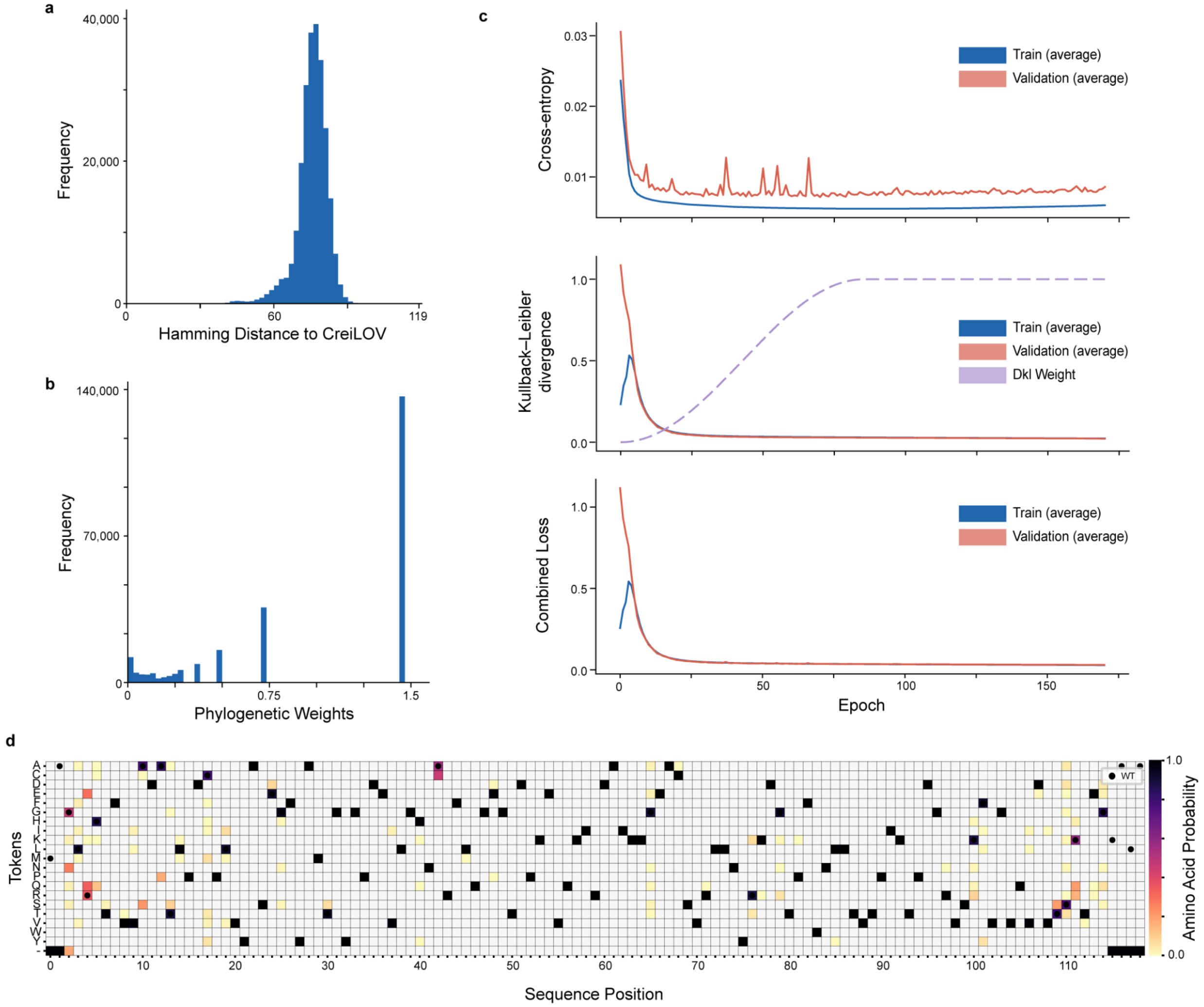


Supplementary Figure 29: VAE pre-training. a, Distribution of sequence identity in curated MSA of natural sequences related to CreiLOV used to train convolutional VAE. b, Distribution of weights for natural sequences in the curated MSA that account for phylogenetic bias in the MSA. c, Training curves for VAE pre-training showing cross-entropy loss, Kullback-Lieber divergence, and combined reconstruction loss (also known as the evidence lower bound) d, Single mutant probabilities for WT (CreiLOV) for VAE. Single mutant probabilities were obtained from CreiLOV sequence input into VAE and taking the softmax of logits for each position of the output sequence generated by VAE.

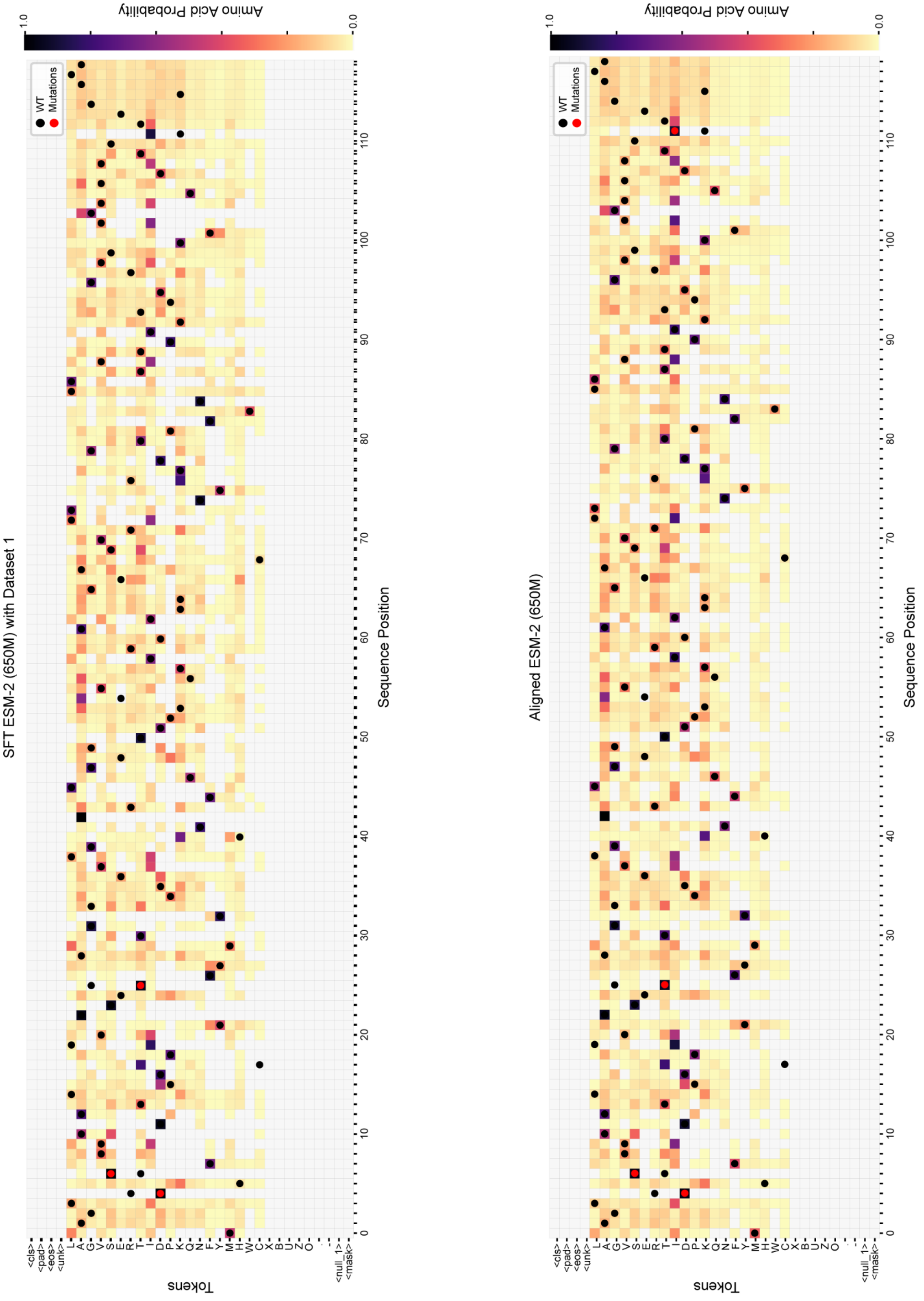


Supplementary Figure 30: Single mutant probabilities for CreiLOV sequence from ESM-2 (650M) after supervised fine-tuning and proximal policy optimization. We indicate high confidence mutations with red circles. Single mutant probabilities were obtained by masking CreiLOV one position at a time and taking the softmax of logits generated by ESM-2 models.

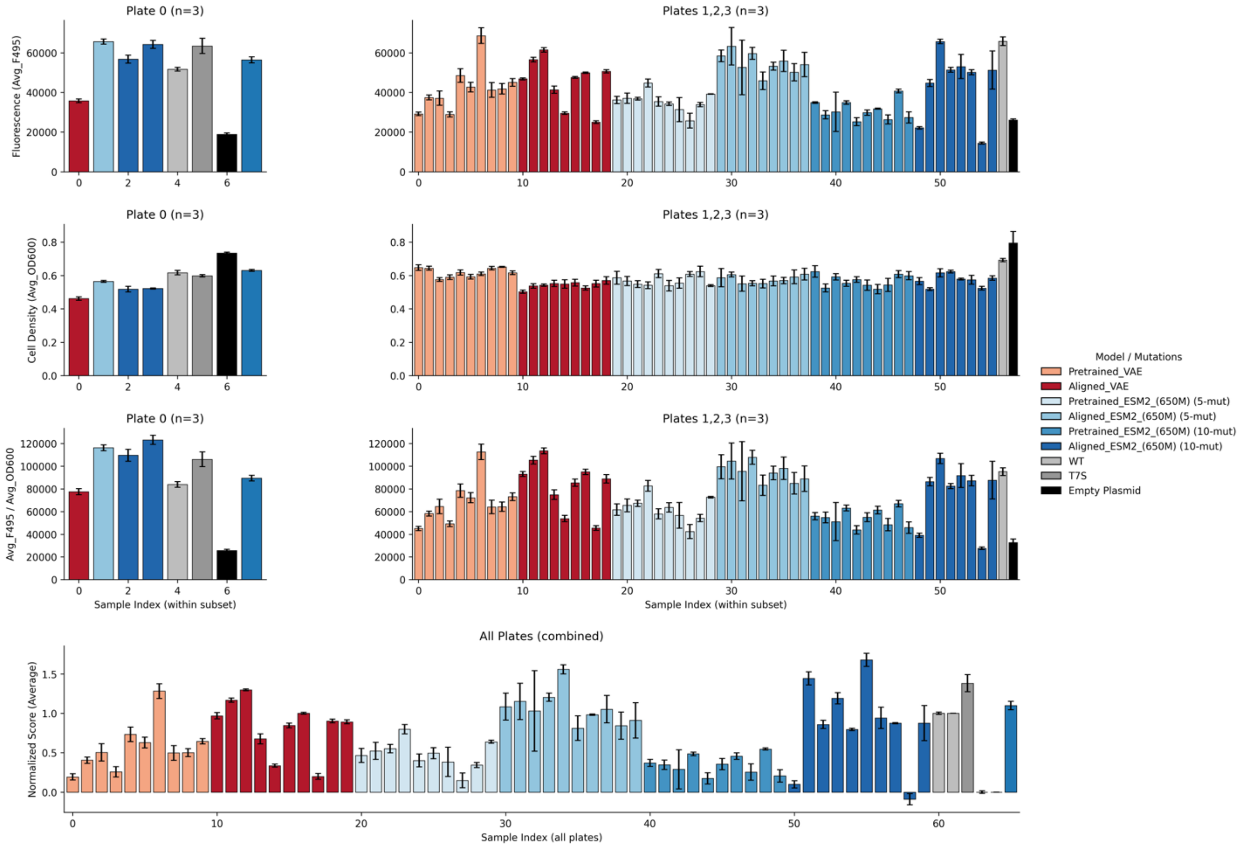


Supplementary Figure 31: Experimental data for variants. We provide the averages and standard deviations for measurements for all sequence variants. We normalize values by empty plasmid and wildtype in the final row to combine plates. For the initial plate, we normalized each replicate by the average empty and wildtype performance. For the following plates, we normalized each replicate by the empty and wildtype performance on that plate. The 5-mutant here is the best 5-mutant present in the CreiLOV DMS dataset.

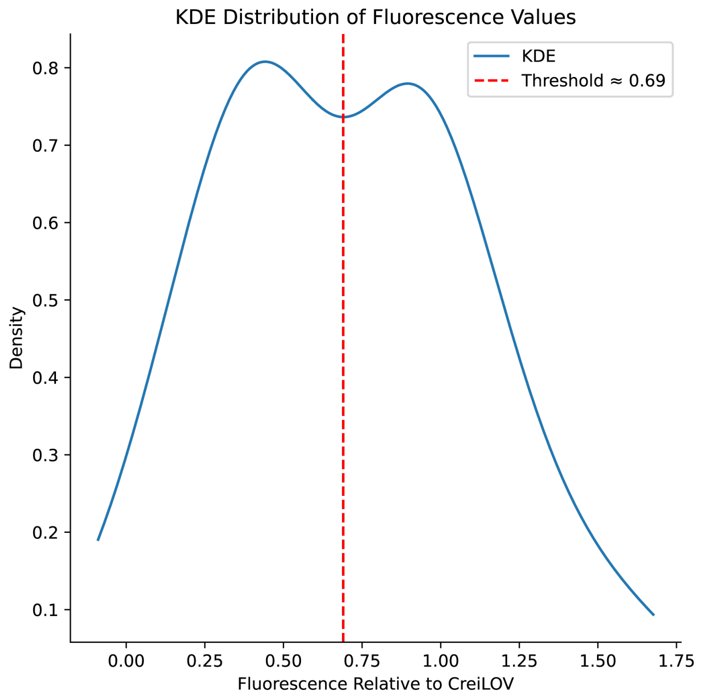


Supplementary Figure 32: Kernel density estimation (KDE) of experimental fluorescence data relative to CreiLOV. The KDE curve reveals a bimodal distribution, indicating the presence of two distinct populations of functional and non-functional variants. A threshold (red dashed line) was determined by locating the minimum between the two highest density peaks, providing a data-driven cutoff to classify sequences based on fluorescence activity.

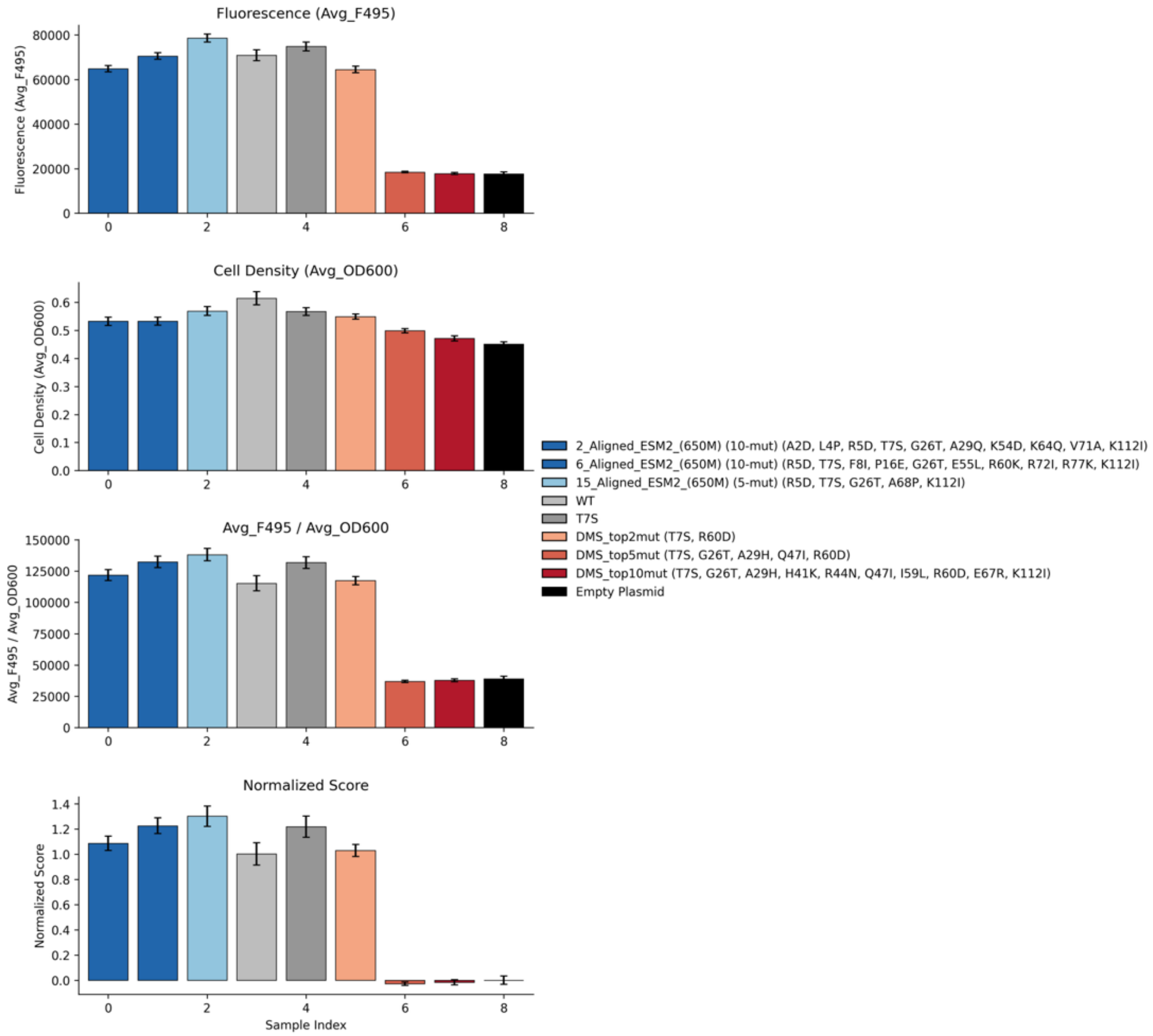


Supplementary Figure 33: *In vivo* fluorescence of sequence variants consisting of the best 2, 5, or 10 single mutant variants. Averages were calculated across 6 replicates.

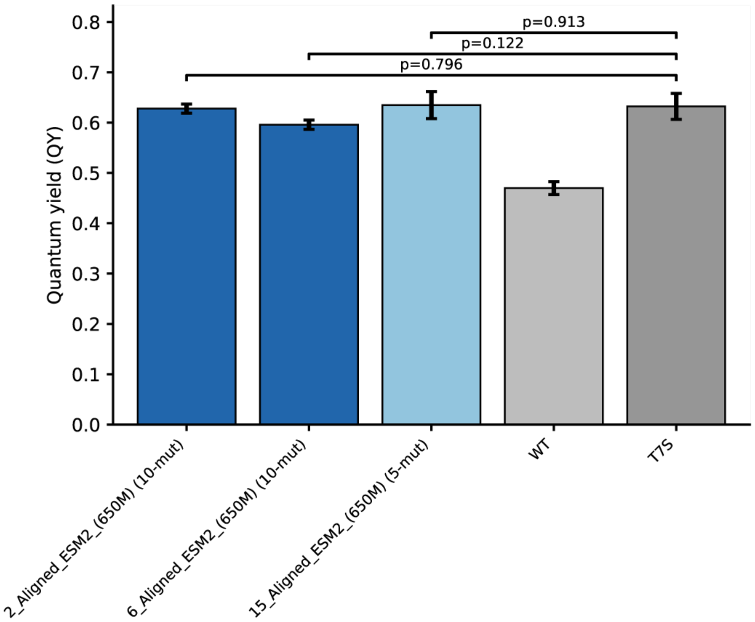


Supplementary Figure 34: Fluorescence quantum yield of CreiLOV variants. The fluorescence quantum yields value obtained for three separate preparations for each variant is plotted. The designed variants exhibited quantum yield value comparable to that of T7S mutant. The value observed for T7S is slightly higher than previously reported value of 0.57. This may be due to differences in experimental condition or the construct design. P values were calculated using Welch's two-sample t-test comparing replicate quantum yield measurements of each aligned variant to T7S.

Supplementary Table 1: DMS Assay Readouts and Evolution

| Protein | DMS assay readout | Rationale |
|---|---|---|
| **Pab1** | Growth selection after endogenous PAB1 gene deletion | Assay captures an essential in vivo function, but the selection is simplified and can over-emphasize stability and RNA-binding. |
| **Bgl3** | Hydrolysis activity via microfluidic-based assay | Assay directly measured enzymatic function, and the DMS landscape was found to strongly correlate with sequence conservation. |
| **Ube4b** | Ubiquitin ligase activity via phage display | Increased activity is not universally beneficial and has been shown to be oncogenic. Natural selection is also constrained by regulation, specificity, and network effects. |
| **avGFP** | Cellular fluorescence via FACS-seq | Assay reflects native output but measured brightness also reflects folding, expression, and maturation in the assay host/conditions. |
| **CreiLOV** | Cellular fluorescence via FACS-seq | Native LOV domains are selected for light sensing/signaling; fluorescence brightness is primarily an engineered reporter objective. |
| **GB1** | IgG-Fc binding via mRNA display | Nature does not select for maximal affinity because GB1 operates in a complex immune environment with trade-offs. |

Proteins were ordered from greatest to least Spearman correlation between DMS assay readout and ESM-2 (650M) evolutionary scores (Supplementary Figure 8).

Supplementary Table 2: Data splits for reward models

| Protein | Mutations | Count | Data split |
|---|---|---|---|
| CreiLOV | 0 | 2 | Training |
| | 1 | 2,204 | Training |
| | 2 | 176 | Training |
| | 3 | 978 | Training |
| | 4 | 3,565 | Training |
| | 5 | 9,603 | 75/25 Val/Test |
| | 6–15 | 151,085 | Not used |
| GB1 | 1 | 1,045 | Training |
| | 2 | 535,039 | 80/15/5 Train/Val/Test |
| avGFP | 1 | 1,114 | Training |
| | 2 | 13,010 | Training |
| | 3 | 12,682 | Training |
| | 4 | 9,759 | Training |
| | 5 | 7,215 | 75/25 Val/Test |
| | 6–15 | 10,243 | Not used |
| Ube4b | 1 | 932 | Training |
| | 2 | 54,507 | Training |
| | 3 | 31,748 | Training |
| | 4 | 8,782 | Training |
| | 5 | 1,903 | 75/25 Val/Test |
| | 6–8 | 425 | Not used |
| Bgl3 | 1 | 3,393 | Training |
| | 2 | 22,465 | Training |
| | 3 | 725 | Training |
| | 4 | 64 | Training |
| | 5 | 6 | 75/25 Val/Test |
| Pab1 | 1 | 1,244 | Training |
| | 2 | 39,608 | 80/15/5 Train/Val/Test |

We split up deep mutational scanning (DMS) datasets by mutational regime for reward model training. For GB1, we randomly sampled half of the double mutants before applying the data split to reduce reward model training time.

Supplementary Table 3: Reward model hyperparameters

| **Hyperparameter** | **CreiLOV** | **GB1** | **avGFP** | **Ube4b** | **Bgl3** |
|---|---|---|---|---|---|
| Loss | MSE | MSE | MSE | MSE | MSE |
| Learning Rate | $1 \times 10^{-6}$ | $1 \times 10^{-6}$ | $1 \times 10^{-6}$ | $1 \times 10^{-6}$ | $1 \times 10^{-6}$ |
| Batch Size | 128 | 128 | 128 | 128 | 128 |
| Epochs | 2000 | 2000 | 2000 | 2000 | 2000 |
| Dropout | 0.1 | 0.1 | 0.1 | 0.1 | 0.1 |
| Patience | 400 | 200 | 400 | 400 | 200 |
| Number of Models | 100 | 3 | 10 | 10 | 10 |
| Hidden Layer Dim. | 400 | 400 | 400 | 400 | 400 |
| Activation | ReLU | ReLU | ReLU | ReLU | ReLU |
| Optimizer | Adam | Adam | Adam | Adam | Adam |
| Embedding Type | One-Hot | One-Hot | One-Hot | One-Hot | One-Hot |

Hyperparameters were selected via grid search for reward models trained with CreiLOV DMS data.

Supplementary Table 4: Reward model correlations

| Correlation | CreiLOV | GB1 | avGFP | Ube4b | Bgl3 | Pab1 |
|---|---|---|---|---|---|---|
| Test Split Pearson's R | 0.940 | 0.972 | 0.774 | 0.573 | 1.0* | 0.931 |
| Test Split Spearman's Rho | 0.933 | 0.972 | 0.691 | 0.471 | 1.0* | 0.923 |

We evaluated reward model accuracy using the test set of variant sequences by calculating Pearson's R and Spearman's Rho between predicted and experimental scores. For Bgl3, there were only 2 sequences in the test set, so the reported correlation values are exact but not statistically meaningful.

Supplementary Table 5: Simulated annealing parameters

| Parameter | CreiLOV | GB1 | avGFP | Ube4b | Bgl3 | Pab1 |
|---|---|---|---|---|---|---|
| Initial temperature | -1.6 | -0.01 | -1.6 | -0.5 | -0.5 | -0.01 |
| Final temperature | -3.1 | -1.5 | -3.1 | -3.1 | -2.25 | -1.8 |
| Number of trials | 100 | 100 | 100 | 100 | 100 | 100 |
| Target mutations per design | 5 | 5 | 5 | 5 | 5 | 5 |
| Number of SA steps | 50,000 | 50,000 | 50,000 | 50,000 | 50,000 | 50,000 |
| Mutations per step | 2 | 2 | 2 | 2 | 2 | 2 |

We performed simulated annealing to generate sequence variants using the parameters listed for each protein. The temperature decreased logarithmically from the initial to final temperature to maintain an initial acceptance probability near 40% for neutral or deleterious mutations, encouraging exploration early in the run, and approaching 0% acceptance by the end of 50,000 steps. It may be necessary to increase the initial acceptance probability for neutral or deleterious mutations for more diverse optimized sequences for performing supervised fine-tuning.

Supplementary Table 6: Supervised fine-tuning hyperparameters

| **Hyperparameter** | **CreiLOV** | **GB1** | **avGFP** | **Ube4b** | **Pab1** | **Bgl3** |
|---|---|---|---|---|---|---|
| Trainable Layers | 25 | 25 | 25 | 25 | 25 | 25 |
| Sequences in dataset | 86 | 24 | 73 | 24 | 43 | 79 |
| Batch Size | 8 | 8 | 8 | 8 | 8 | 8 |
| Epochs | 1 | 1 | 1 | 1 | 1 | 1 |
| Optimizer | Adam | Adam | Adam | Adam | Adam | Adam |
| Base learning rate | 5.11e−3 | 5.11e−3 | 5.11e−3 | 5.11e−3 | 1.02e−2 | 5.11e−3 |
| Learning rate multiplier | 8.90e−1 | 8.90e−1 | 8.90e−1 | 8.90e−1 | 8.90e−1 | 8.90e−1 |
| Weight decay | 3.51e−3 | 3.51e−3 | 3.51e−2 | 3.51e−3 | 3.51e−3 | 3.51e−3 |
| LR scheduler: cosine annealing | True | True | True | True | True | True |
| LR scheduler: warm restart | True | True | True | True | True | True |
| $T_0$ (warm restart period) | 10 | 10 | 10 | 10 | 10 | 10 |
| $T_{mult}$ (multiplier) | 1 | 1 | 1 | 1 | 1 | 1 |
| Gradient Clipping Threshold | 3 | 3 | 3 | 3 | 3 | 3 |

Hyperparameters were selected by Optuna after 1,000 trials of hyperparameter optimization with a Tree-Parzen Estimator (TPE) sampler and a custom PyTorch callback to terminate trials if model parameters became non-finite (i.e., NaN or infinity) after each min-batch. For other proteins, we altered the learning rate or weight decay to increase generative performance.

Supplementary Table 7: Proximal policy optimization hyperparameters with ESM-2

| **Hyperparameter** | **CreiLOV** | **GB1** | **avGFP** | **Ube4b** | **Pab1** | **Bgl3** |
|---|---|---|---|---|---|---|
| Initial trainable layers | 27 | 27 | 27 | 27 | 27 | 27 |
| Layers to unfreeze/epoch | 69 | 69 | 69 | 69 | 69 | 69 |
| Max trainable layers | 82 | 82 | 82 | 82 | 82 | 82 |
| Initial Batch size | 1 | 1 | 1 | 1 | 1 | 1 |
| Increment to batch size/epoch | 1 | 1 | 1 | 1 | 1 | 1 |
| Max batch size | 10 | 10 | 10 | 10 | 10 | 10 |
| Epochs | 2 | 10 | 6 | 2 | 5 | 2 |
| Mutations | 15 | 5 | 6 | 10 | 10 | 10 |
| High-confidence probability threshold | 0.90 | 0.90 | 0.90 | 0.90 | 0.95 | 0.95 |
| Cumulative probability threshold | 0.22 | 0.25 | 0.20 | 0.22 | 0.25 | 0.20 |
| Epsilon | 1.74e−1 | 1.74e−1 | 1.74e−1 | 1.74e−1 | 1.74e−1 | 1.74e−1 |
| Initial $D_{KL}$ weight | 1.00e−8 | 1.00e−8 | 1.00e−8 | 1.00e−8 | 1.00e−8 | 1.00e−8 |
| $D_{KL}$ weight | 1.00e−7 | 1.00e−7 | 1.00e−7 | 1.00e−7 | 1.00e−7 | 1.00e−7 |
| Pairwise Hamming distance weight | 1.00e−6 | 1.00e−6 | 1.00e−6 | 1.00e−6 | 1.00e−6 | 1.00e−6 |
| Optimizer | Adam | Adam | Adam | Adam | Adam | Adam |
| Base learning rate | 8.66e−3 | 8.66e−3 | 8.66e−3 | 8.66e−3 | 8.66e−3 | 8.66e−3 |
| Learning rate multiplier | 0.885 | 0.885 | 0.885 | 0.885 | 0.885 | 0.885 |
| Weight decay | 9.95e−3 | 9.95e−3 | 9.95e−3 | 9.95e−3 | 9.95e−3 | 9.95e−3 |
| LR scheduler: cosine annealing | True | True | True | True | True | True |
| LR scheduler: warm restart | True | True | True | True | True | True |
| Grad clip threshold | 6.824 | 6.824 | 6.824 | 6.824 | 6.824 | 6.824 |
| Exponential moving average decay | 0.8 | 0.8 | 0.8 | 0.8 | 0.8 | 0.8 |

Hyperparameters for aligning ESM-2 were identified with Optuna using reward models trained with CreiLOV DMS data. For other proteins, we varied the number of epochs, mutations, and probability thresholds for sampling to increase generative performance.

Supplementary Table 8: VAE pre-training hyperparameters

| **VAE Hyperparameters** | |
|---|---|
| Learning Rate | $1 \times 10^{-4}$ |
| Batch Size | 32 |
| Epochs | 1000 |
| Patience | 100 |
| Weighted $D_{KL}$ | Cyclical Annealing Schedule |
| Number of Cycles | 1 |
| Embedding Type | One-Hot |
| 1st Convolutional Layer | 21 input channels, 21×16 output channels |
| 2nd Convolutional Layer | 21×16 input channels, 21 output channels |
| Kernel size | 17 |
| Padding | 1 |
| Fully Connected Layer Dim. | 400 |
| Latent Space Dim. | 64 |
| Optimizer | Adam |
| Activation | ReLU for fully-connected layers |
| Activation | LeakyReLU for convolutional layers |

Hyperparameters were selected via grid search of 673 combinations. ReLU activations were applied to fully connected layers. LeakyReLU activations were applied to convolutional layers.

Supplementary Table 9: Proximal policy optimization hyperparameters with VAE

| **Hyperparameter** | **CreiLOV** |
|---|---|
| Initial trainable layers | 6 |
| Layers to unfreeze/epoch | 0 |
| Max trainable layers | 6 |
| Initial batch size | 29 |
| Increment to batch size/epoch | 9 |
| Max batch size | 64 |
| Epochs | 27 |
| Target mutations | 5 |
| Amino acid sampling | Max Likelihood for Position |
| Iterations | 4 |
| Epsilon | 0.1826 |
| Initial $D_{KL}$ weight | 1.00e−8 |
| $D_{KL}$ weight | 1.00e−7 |
| Pairwise Hamming distance weight | 89.82 |
| Optimizer | Adam |
| Base learning rate | 6.52e−4 |
| Learning rate multiplier | 0.8821 |
| Learning rate multiplier factor | 0.9718 |
| Weight decay | 2.99e−6 |
| LR scheduler: cosine annealing | True |
| LR scheduler: warm restart | True |
| Grad clip threshold | 2.7783 |
| Grad clip threshold factor/epoch | 2 |
| Exponential moving average decay | 0.992 |

Hyperparameters were identified with Optuna with reward models trained with CreiLOV DMS data.

Supplementary Table 10: Curation of multiple sequence alignments for EVE training

| Protein | Domain-level threshold | Jackhmmer iterations | Min column coverage | Min sequence coverage | Sequence filter | Sequence-level threshold | MSA length (N) | Neff |
|---|---|---|---|---|---|---|---|---|
| avGFP | 0.5 | 5 | 70 | 50 | – | 0.5 | 1087 | 151.196 |
| Bgl3 | 0.5 | 5 | 70 | 50 | 90 | 0.5 | 17139 | 8692.459 |
| CreiLOV | 0.3 | 5 | 70 | 50 | 90 | 0.3 | 43374 | 36700.867 |
| GB1 | 0.3 | 5 | 70 | 50 | 95 | 0.3 | 12110 | 1554.414 |
| Pab1 | 0.4 | 4 | 70 | 70 | 80 | 0.4 | 5742 | 5622.467 |
| Ube4b | 0.3 | 5 | 70 | 50 | 85 | 0.3 | 10707 | 6538.152 |

Summary of the MSA retrieval and filtering parameters used to curate UniRef100 alignments for each benchmark protein to training family-specific EVE models, including domain- and sequence-level coverage thresholds, number of Jackhmmer iterations, sequence filtering criteria, and the resulting MSA depth (number of sequences, N) and effective number of sequences (Neff).

Supplementary Table 11: Win rate of sequence variants after PPO

| Model | Summary Statistics | | | | Win Rate | | |
|---|---|---|---|---|---|---|---|
| | Mean | Median | Max | Min | vs. VAE | vs. ESM-2 (650M) | vs. Aligned ESM-2 (650M) |
| VAE | 4.039 | 4.038 | 4.095 | 3.967 | - | 0.540 | 0.0000 |
| Aligned VAE | 4.086 | 4.094 | 4.115 | 4.013 | 0.890 | 0.920 | 0.000 |
| ESM-2 (8M) | 4.029 | 4.039 | 4.126 | 3.863 | 0.510 | 0.570 | 0.000 |
| SFT ESM-2 (8M) | 4.170 | 4.174 | 4.206 | 4.050 | 1.000 | 1.000 | 0.280 |
| Aligned ESM-2 (8M) | 4.169 | 4.177 | 4.211 | 4.087 | 1.000 | 1.000 | 0.330 |
| ESM-2 (35M) | 4.018 | 4.022 | 4.103 | 3.921 | 0.260 | 0.370 | 0.000 |
| SFT ESM-2 (35M) | 4.052 | 4.052 | 4.052 | 4.052 | 0.840 | 0.910 | 0.000 |
| Aligned ESM-2 (35M) | 4.046 | 4.046 | 4.046 | 4.046 | 0.720 | 0.810 | 0.000 |
| ESM-2 (150M) | 4.016 | 4.025 | 4.116 | 3.875 | 0.270 | 0.370 | 0.000 |
| SFT ESM-2 (150M) | 4.172 | 4.174 | 4.203 | 4.121 | 1.000 | 1.000 | 0.260 |
| Aligned ESM-2 (150M) | 4.170 | 4.176 | 4.200 | 4.113 | 1.000 | 1.000 | 0.240 |
| ESM-2 (650M) | 4.030 | 4.035 | 4.123 | 3.962 | 0.460 | - | 0.000 |
| SFT ESM-2 (650M) | 4.184 | 4.188 | 4.210 | 4.133 | 1.000 | 1.000 | - |
| Aligned ESM-2 (650M) | 4.199 | 4.202 | 4.215 | 4.164 | 1.000 | 1.000 | 0.840 |

We sample 100 sequences from each model to calculate summary statistics and win rate. Interestingly, the median predicted log fluorescence for sequences generated from aligned ESM2 (650M) was greater than the max predicted log fluorescence for sequences generated via simulated annealing. 92 of the first 100 generated sequences from aligned ESM-2 (650M) were unique.
